# H3K4me3 enrichment defines neuronal age, while a youthful H3K27ac signature is recapitulated in aged neurons

**DOI:** 10.1101/2021.11.11.467877

**Authors:** Katherine A. Giles, Andrew J. Phipps, Jake M. Cashion, Shannon N. Huskins, Timothy R. Mercer, Mark D. Robinson, Adele Woodhouse, Phillippa C. Taberlay

## Abstract

Neurons live for the lifespan of the individual and underlie our ability for lifelong learning and memory. However, aging alters neuron morphology and function resulting in age-related cognitive decline. It is well established that epigenetic alterations are essential for learning and memory, yet few neuron-specific genome-wide epigenetic maps exist into old age. Comprehensive mapping of H3K4me3 and H3K27ac in mouse neurons across lifespan revealed plastic H3K4me3 marking that differentiates neuronal age linked to known characteristics of cellular and neuronal aging. We determined that neurons in old age recapitulate the H3K27ac enrichment at promoters, enhancers and super enhancers from young adult neurons, likely representing a re-activation of pathways to maintain neuronal output. Finally, this study identified new characteristics of neuronal aging, including altered rDNA regulation and epigenetic regulatory mechanisms. Collectively, these findings indicate a key role for epigenetic regulation in neurons, that is inextricably linked with aging.

## INTRODUCTION

Aging is characterised by the gradual loss of physiological integrity resulting in the impaired function of bodily systems. In the central nervous system, aging is associated with variable cognitive decline in healthy individuals. The activity of neurons in the brain underlies cognitive function, and like many bodily systems, aging of the central nervous system is linked to functional and structural vicissitudes in neurons^1,2^. Neurons are one of the few cell types in the body that can subsist for the entire lifespan. Although neurons exit the cell cycle, functional plasticity is retained to enable lifelong learning and memory. Changes in epigenetic marking are inextricably linked with learning and memory, as well as with age-related cognitive decline^3–5^.

The epigenome is at the interface of genes and the environment, enabling dynamic regulation of gene expression in each cell. The ‘epigenome’ includes DNA methylation and histone modifications, as well as the physical organisation of DNA inside the nucleus, which dynamically regulates gene expression^6^. The combination of epigenetic elements determines whether a genomic region is active, repressed or silenced. Yet like all bodily systems, age-related alterations in the epigenome occur in cells linked to a decline in function^7^. Added to this, the long lifespan of neurons may enable epigenetic “errors” to accumulate over time.

Several studies have established that age-associated epigenetic alterations occur in the brain^8–11^. Histone modifications correlate with age-associated memory deficits, while increased histone acetylation and inhibition of histone de-acetylation pathways can reverse age-associated cognitive deficits^12–15^. However, the brain comprises a complex mixture of cell types, and each cell type in the brain has a unique epigenetic signature^8,16–19^. Furthermore, age-related changes to the epigenome differ between neurons and glia^8,10,16,17^. In 2013, two seminal papers describing epigenetic signatures in neurons across lifespan were published^16,20^. Human neurons from the prefrontal cortex exhibited dynamic histone 3 lysine 4 tri-methylation (H3K4me3) marking from prenatal development to adulthood, which remained relatively stable into old age^20^. However, in mouse hippocampal neurons genome-wide analysis revealed histone 2 lysine 12 acetylation (H3K12ac) depletion with age^8^. Few genome-wide epigenetic maps of purified neurons exist from adulthood into old age^8,10,16,20^, and we are just beginning to unravel how the multi-layered epigenetic landscape of neurons changes across a lifetime.

Here we have produced comprehensive maps of H3K4me3 and histone 3 lysine 27 acetylation (H3K27ac) dynamics in purified neurons across lifespan in C57/BL6 mice. These maps reveal dynamic H3K4me3 marking that differentiates between neurons of different ages. We also identify that neurons in old age recapitulate the H3K27ac enrichment at promoters, enhancers and super enhancers that were present in young adult neurons. Furthermore, this work has discovered epigenetic regulation and rDNA regulation as new characteristics of neuronal aging.

## RESULTS

### Generation of epigenome maps for H3K4me3 and H3K27ac during neuron aging

To date, there are few epigenetic maps of neurons from adulthood into old age^8,10,16,20^. To address the need for neuronal epigenetic information we optimised a protocol using fluorescence activated nuclei sorting (FANS) protocol to purify NeuN^+^ (RNA binding protein, fox-1 homolog; *Rbfox3* positive) neuronal nuclei from the forebrains of C57/BL6 mice (Extended Data Figure 1A-B). We applied the FANS neuronal nuclei purification technique to the forebrains of C57/BL6 mice at 3 months (young adult), 6 months (mature adult), 12 months (middle age) and 24 months of age (old age; Figure 1A-B). Post-purification, the chromatin of the neuronal nuclei was immediately cross-linked then subject to chromatin immunoprecipitation and sequencing (ChIP-seq; Figure 1C-D) to profile the dynamics of H3K4me3 and H3K27ac across lifespan. Both H3K4me3 and H3K27ac are associated with the promoters of actively expressed genes^6,21^, and are key predictors of age-related transcriptional change in the brain^11,22^. In addition, H3K27ac is associated with active enhancers and super enhancers, which are linked to cell identity^6,21^. Given these histone modifications have been extensively characterised as features of DNA regulatory elements they were ideal to characterise the neuronal aging epigenome.

**Figure 1.**
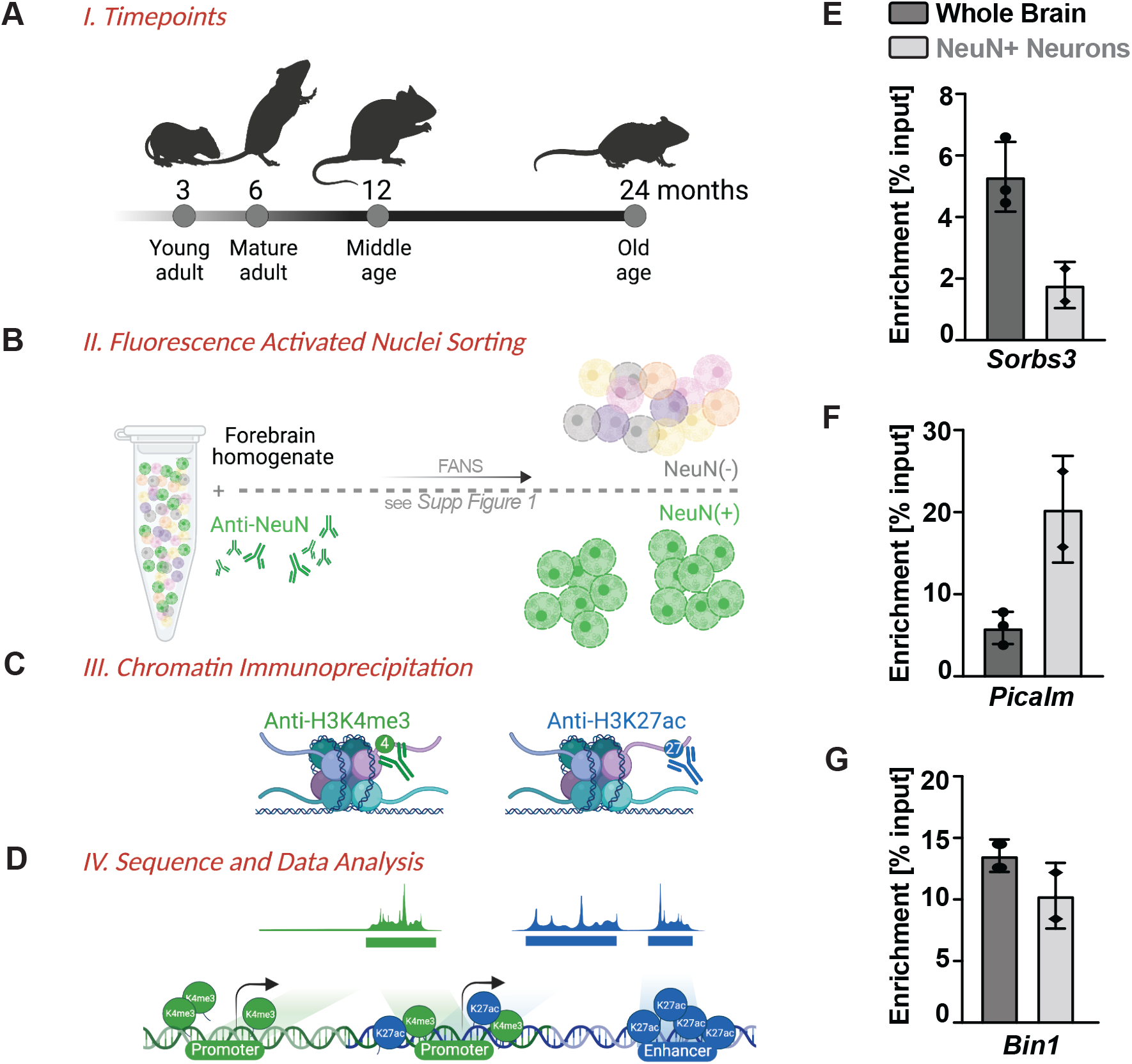
Genome-wide epigenome maps of aging neurons. **A-D)** Schematic of experimental design and analysis for this study, including (**A**) Timeline of mouse lifespan and data points collected for this study, (**B**) FANS collection using NeuN+ cells, **C**) Chromatin immunoprecipitation and (**D**) sequencing. Created with BioRender.com. **E-G)** ChIP-qPCR targeting H3K27ac of mouse whole brain (n=2-3) and NeuN+ purified neurons (n=2) for promoters *Sorbs3* (**E**), *Picalm* (**F**) and *Bin1* (**G**). Enrichment is ChIP-qPCR signal over IgG and over total input.

To first illustrate that we can enrich for a neuronal epigenetic signature we performed ChIP-qPCR for H3K27ac and compared our purified neuronal nuclei and to whole brain lysates. Differences in H3K27ac enrichment at the promoter of *Sorbs3* were detected, with enrichment in the whole brain sample (Figure 1E). Conversely, there was higher enrichment of H3K4me3 at the *Picalm* promoter in neuronal nuclei compared to the whole brain (Figure 1F). Meanwhile, the promoter region of *Bin1* showed similar enrichment in whole brain and the neuronal sample (Figure 1G). These data demonstrate that purified neurons have a unique neuronal epigenetic signature that is masked when examining whole brain lysate.

We then examined our genome-wide sequencing data from purified neurons at the promoters of the neuronal specific genes *Syt1* and *Syp*. As expected, there was a robust signal for H3K4me3 at the promoters of these genes (Figure 2A). However, H3K27ac was more plastic at these genes across the neuronal lifespan. Conversely, at genes that are expressed in glial cells and not in neurons, for example *Gfap* and *Aif1*, no signal was detected for either H3K4me3 or H3K27ac (Extended Data Figure 2A). We also examined the regions analysed in the ChIP-qPCR and found that H3K27ac was also enriched at the *Picalm1* and *Bin1* promoters in the genome wide dataset (Extended Data Figure 2B). Moreover, there was no enrichment at the *Sorbs3* promoter, which we had shown was more highly enriched in the whole brain sample (Extended Data Figure 2B). Therefore, our data shows the expected pattern of H3K4me3 and H3K27ac enrichment at neuronal marker genes and supports our preparation of purified neurons.

**Figure 2.**
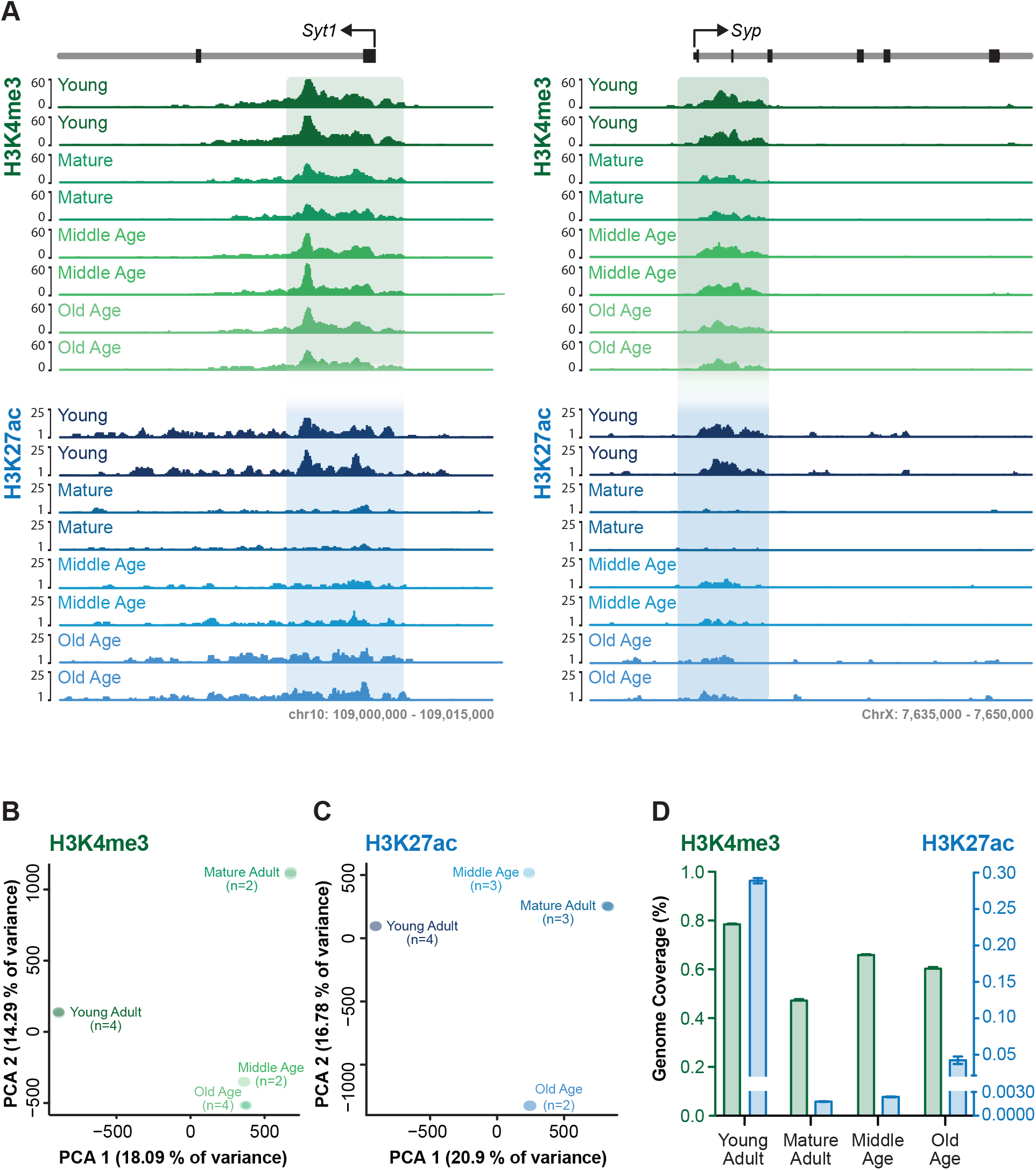
Enrichment of H3K4me3 and H327ac in neurons. **A)** Example regions of H3K4me3 and H3K27ac neuronal ChIP-seq signal at neuron specific genes *Syt1* and *Syp,* displaying two representative tracks for each age. **B-C)** PCA plots for genome wide ChIP-seq signal of (B) H3K4me3 and (C) H3K27ac in neurons with the first dimension plotted on the x-axis and second dimension on the y-axis. Plots show clustering of the samples by replicate and separation of samples by age. **D)** Percent in base pairs of the genome covered by H3K4me3 and H3K27ac in neurons for each age. The left y-axis and green bars display H3K4me3 data, while the right y-axis and blue bars represent H3K27ac.

### H3K4me3 and H3K27ac enrichment is dynamic during neuronal aging

We examined our data for the similarities and variations in H3K4me3 and H3K27ac enrichment in neurons with age. To compare the genome wide features of our data we calculated the ChIP-seq signal enrichment in each sample across the genome. Pearson correlation analysis comparing each sample within each histone mark revealed that the correlations were the strongest between samples from the same time point, as expected (Extended Data Figure 3A-B). Similarly for Principal Component Analysis (PCA), samples for each time point clustered strongly together (Figure 2B-2C). In the PCA for H3K4me3, the young adult neurons separated from the other time points on the first dimension, and in the second dimension, the mature adult neurons were significantly separated from the middle and old age neurons (Figure 2B). A comparable pattern was observed for H3K27ac, however the middle age neurons clustered closer to the mature adult neurons on the second dimension (Figure 2C). Together, these analyses indicate that a leading source of variation in our datasets is related to age.

We next considered whether the proportion of the genome marked by these histone modifications changes during neuronal aging. We assessed the genome coverage (in base pairs) using MACS2 peak calling and found both histone marks varied by ∼0.3% across the time course. H3K4me3 ranged from ∼0.5-0.8% and H3K27ac ranged from ∼0.001-0.3% (Figure 2D). The young adult mice exhibited the highest genome coverage for both histone marks, which reduced in mature adults. Whilst H3K4me3 coverage is reacquired in middle age and then retained in old age, H3K27ac is not regained until old age. This suggests that while globally these marks are, in part, re-established in aging neurons, this occurs independently for individual marks.

### H3K4me3 is redistributed at promoters across neuronal lifespan

To determine which genome regions exhibited variation in H3K4me3 enrichment, we next performed chronological pairwise differential analysis. Using the genome-wide ChIP-seq signal and a 150bp sliding window we compared 1) young and mature adult neurons, 2) mature and middle age neurons and 3) middle and old age neurons. Differential regions required a minimum of 1000bp between them, and reported as up or down based on the largest change that occurred within the region. We detected 27,769 H3K4me3 regions that changed between young and mature adult neurons, and strikingly the vast majority (94%) were down regulated (Figure 3A). Fewer changes were observed between mature adult and middle age neurons (18,825) and between middle and old age (22,162; Figure 3B-C). In contrast to younger neurons, most of the regions detected here were up regulated with increased age (74% and 63% respectively; Figure 3B-C). This supports the global pattern of H3K4me3 we identified across aging (Figure 2D).

**Figure 3.**
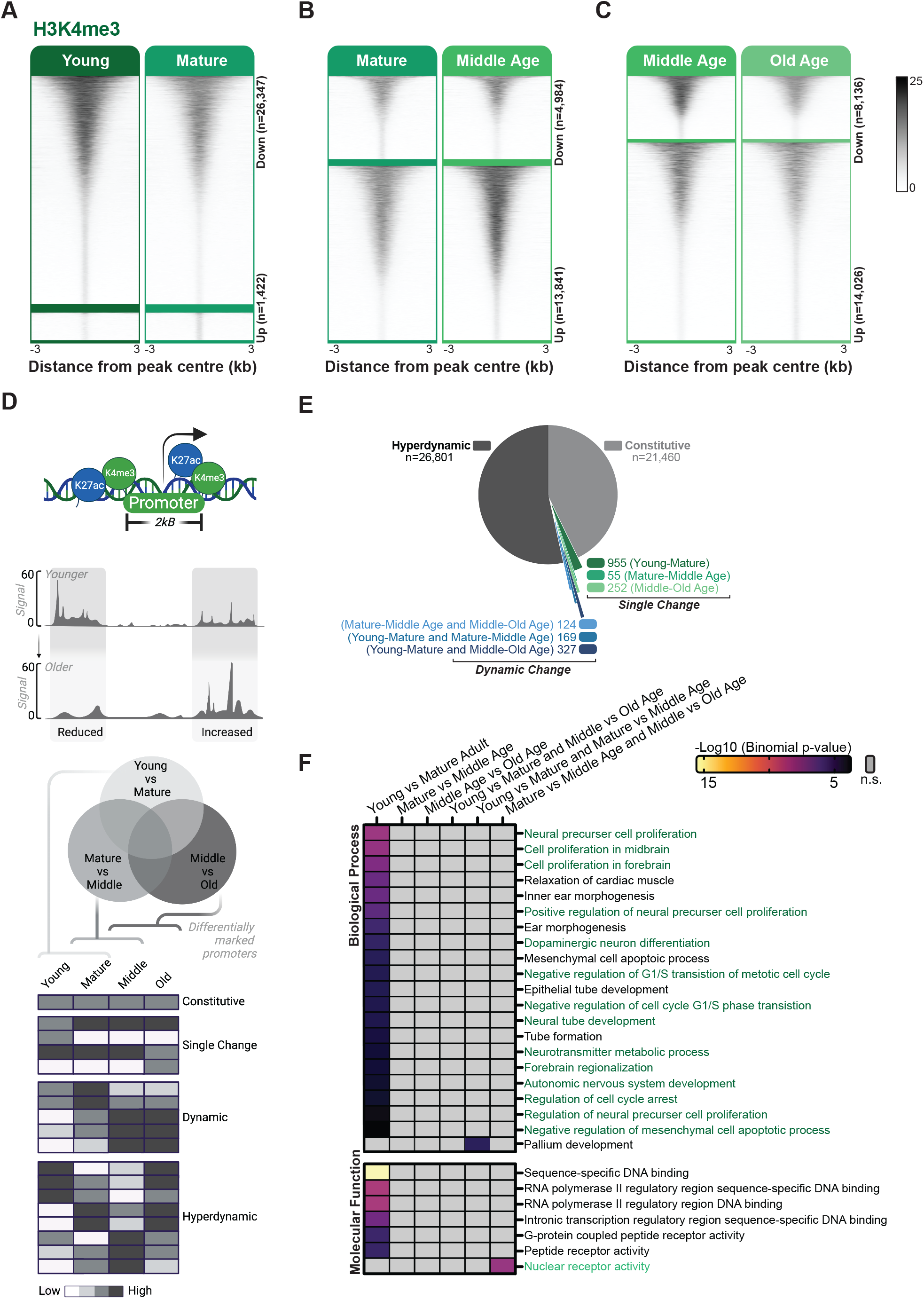
H3K4me3 at Promoters. **A-C)** Heatmaps of H3K4me3 signal at differential regions for (**A**) young versus mature adults, (**B**) mature versus middle age adults and (**C**) middle versus old age adults. Signal is positioned on the centre +/- 3 kb of each differential region and sorted in descending order. **D)** Schematic of analysis to determine the categories of promoter groups. After the overlap of annotated promoters and differential ChIP-seq regions, examples are shown of promoters that are either constitutive (no change including no signal), single change (changes once at any point in the time course), dynamic (change twice) or hyperdynamic (changes at each point across the time course). Created with BioRender.com. **E)** The percentage of promoters within each category based on H3K4me3 signal. Single and dynamic changes are further classified based on which time point the change occurred. **F)** Significant gene ontology biological process terms for a change in H3K4me3 with either a signal or dynamic change. Terms associated with neuronal development, the cell cycle or proliferation are denoted in green.

The most well-known function of H3K4me3 is to mark the promoters of actively transcribed genes. Therefore, we interrogated our data to determine the impact of H3K4me3 at promoters during neuronal aging. We categorised each promoter for H3K4me3 signal as either; 1) constitutive, where no statistically significant change in signal was detected, including promoters where no signal was detected across the entire dataset; 2) single change, where there is a significant change in signal at one time point and that change persists for the remainder of the lifespan; 3) dynamic, where the promoter signal changes twice across aging or 4) hyperdynamic, where a change in signal at the promoter was detected at each time point across the adult mouse lifespan (Figure 3D). In this analysis, we observed 21,460 H3K4me3 constitutively marked promoters (Figure 3E). There were 1,262 promoters exhibiting a single change, with the majority of these (955; 76%) occurring between young and mature adults (Figure 3E). We also detected 620 regions with dynamic changes that were spread across the lifespan (Figure 3E). Overwhelmingly, the promoters with a change in H3K4me3 were hyperdynamic, with 26,801 regions included in this group (Figure 3E).

We next performed a gene ontology (GO) analysis using GREAT to identify gene pathways and regulatory networks with a change in H3K4me3 in aging neurons. For promoters with a single change in H3K4me3 between young and mature adults, we found a significant enrichment of GO terms relating to brain development and cell proliferation (Figure 3F). This suggests that neuron maturation, differentiation and cell cycle processes are still occurring between neurons in young and mature adulthood. In addition, the term ‘Pallium development’ was enriched for dynamic promoters with a change between young and mature, and mature and middle age neurons (Figure 3F). The GO term ‘nuclear receptor activity’ was enriched for dynamic promoters with a change in H3K4me3 between middle age and mature neurons, and between mature and old age neurons, suggesting hormone activity is important during the aging process (Figure 3F). The remaining single change and dynamic groups were not enriched for any GO terms. The hyperdynamic group was linked with numerous GO terms, which warranted further investigation of this group to determine how these pathways were affected across the mouse lifespan.

To probe the hyperdynamic group further we performed hierarchical clustering of the H3K4me3 signal at the 26,801 promoters. We identified four clusters of H3K4me3 signal enrichment each peaking at a different phase of adulthood (Figure 4A), with multiple GO terms associated with each cluster (Figure 4B and Extended Data Figure 4A, 5A). In the cluster that had the highest H3K4me3 enrichment in young adults and the lowest in old age (cluster 3), the most enriched pathways were relating to telomeres and mitochondrial function. Additionally, this cluster was enriched for spliceosome and DNA repair pathways (Figure 4B and Extended Data Figure 4A, 5A). Promoters that had most H3K4me3 enrichment in mature adult neurons and the least in aged neurons (cluster 2) were also associated with mitochondrial function, although these pathways were less significant than in cluster 3 (Figure 4B and Extended Data Figure 4A). In the cluster characterised by promoters that had the highest H3K4me3 marking in middle age and the least in old age (cluster 1), linked GO pathways included base-excision repair, rRNA processing, strand elongation and synaptic vesicle transport (Extended Data Figure 4A, 5A). Where promoters were highly marked with H3K4me3 in aged neurons with some enrichment in young neurons (cluster 4), we observed associations with cellular response to dsRNA and gene silencing by RNA (Extended Data Figure 4A, 5A). We also detected pathways enriched across multiple time points, which notably were all more statistically significant in younger neurons and the significance decreases with age. These include pathways known to be altered in neurons with age including DNA replication-dependent nucleosome assembly, protein folding, ncRNA processing and mitochondrial pathways. Moreover, we identified a novel pathway associated with neuronal aging; chromatin silencing at rDNA (Figure 4B and Extended Data Figure 4A). Together, this shows that the epigenomic signature of H3K4me3 is plastic and enriches for different pathways over neuronal lifespan.

**Figure 4.**
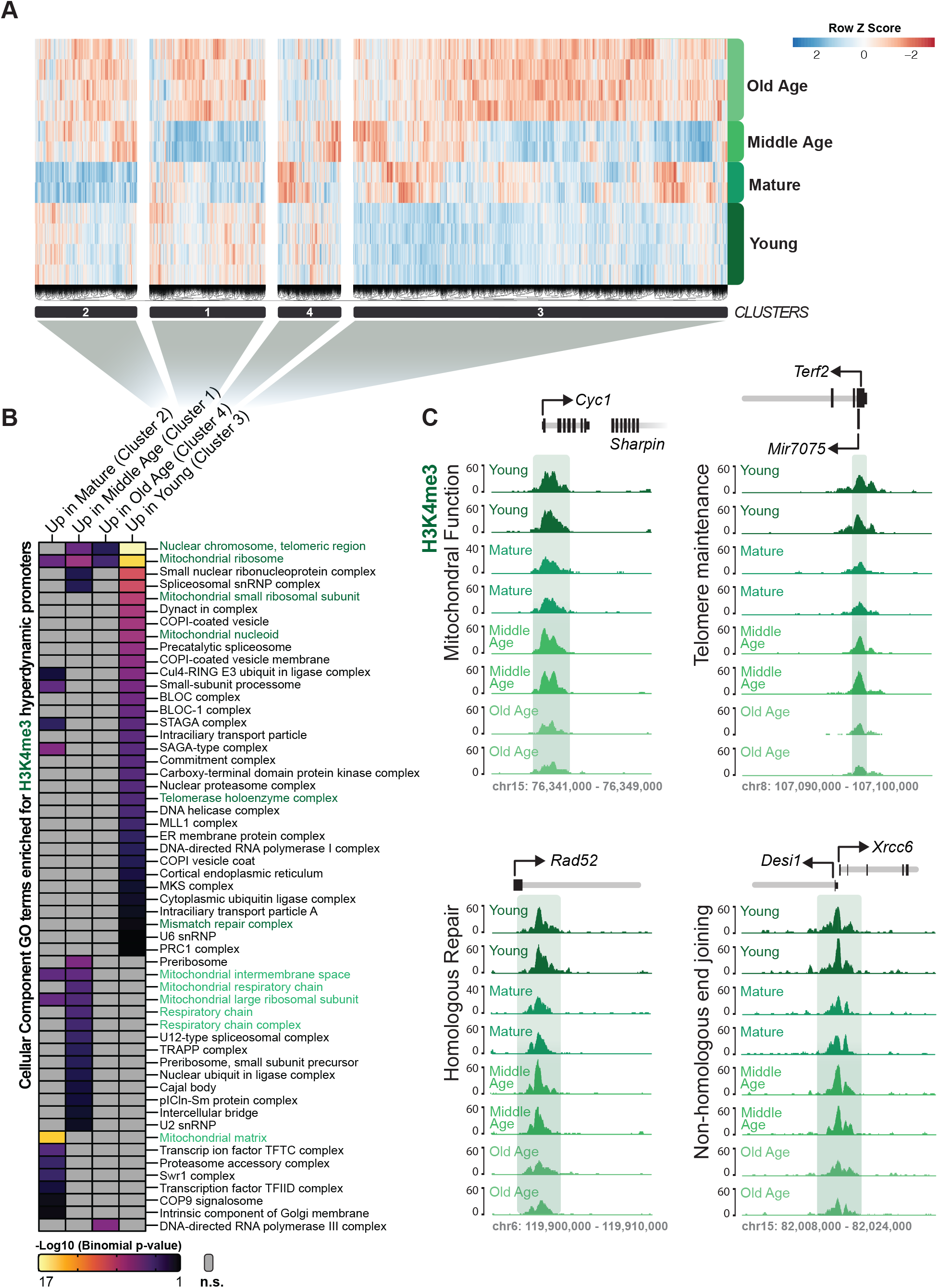
H3K4me3 at Hyperdynamic Promoters. **A)** Heatmap of normalised (row Z score) ChIP-seq signal of H3K4me3 at hyperdynamic promoters. Promoters are divided into four groups by hierarchical clustering and numbered based on position in the dendrogram. **B)** Cellular component gene ontology terms associated with hyperdynamic promoters. Terms relating to telomeres, mitochondrial function or DNA repair are denoted in green. **C)** Example regions of H3K4me3 signal at *Cyc1* (mitochondrial function), *Terf2* (telomere maintenance), *Rad52*and *Xrcc6* (DNA repair). Each region displays two representative tracks for each age.

Analysis of gene expression has identified the down regulation of telomere maintenance, mitochondrial function and DNA repair genes as known hallmarks of cellular aging^23,24^. Since we similarly identified pathways related to these in cluster 3 of the hyperdynamic promoters (highest in young, lowest in old age cluster), we examined the gene hits in these pathways more closely. Examples of reduced H3K4me3 signal with aging can be seen at the promoters for *Cyc1* and *Cycs* for mitochondrial function (Figure 4C and Extended Data Figure 5B), *Terf1* and *Terf2* for telomere maintenance (Figure 4C and Extended Data Figure 5B), and several DNA repair genes such as *Xrcc1, Lig4, Pold3, Xrcc6, Rad52, Brca2* (Figure 4C and Extended Data Figure 5B).

Together, this analysis of dynamic epigenetic modifications identifies known traits of cellular and neuronal aging, such as alterations in DNA repair, protein folding, telomere maintenance, mitochondrial function, non-coding RNA regulation and synaptic function. Furthermore, these data highlight the alterations of nucleosome assembly in aging neurons and support rDNA chromatin regulation being a hallmark of neuronal aging.

### Age-related H3K27ac plasticity at promoters demonstrated a recapitulation of the early adulthood profile in aged neurons

We performed the same differential analysis for H3K27ac in the aging neurons, where we observed fewer changes than for H3K4me3, but detected a more dynamic enrichment pattern. Between young and mature adults, there was a large decrease in H3K27ac signal at promoters where 99% of the 11,867 differential regions were down regulated (Figure 5A). Far fewer changes occurred between mature adult and middle aged neurons (2,595), with 99% of the promoters demonstrating increased H3K27ac marking (Figure 5B). Between middle age and old age, there were 9,516 differentially marked promoters detected in neurons, with 88% of them exhibiting increased levels of H3K27ac (Figure 5C). Thus, as seen with the global coverage levels (Figure 2D), the neuronal H3K27ac landscape demonstrated a dramatic loss between young and mature adults, with H3K27ac enrichment progressively increasing until old age.

**Figure 5.**
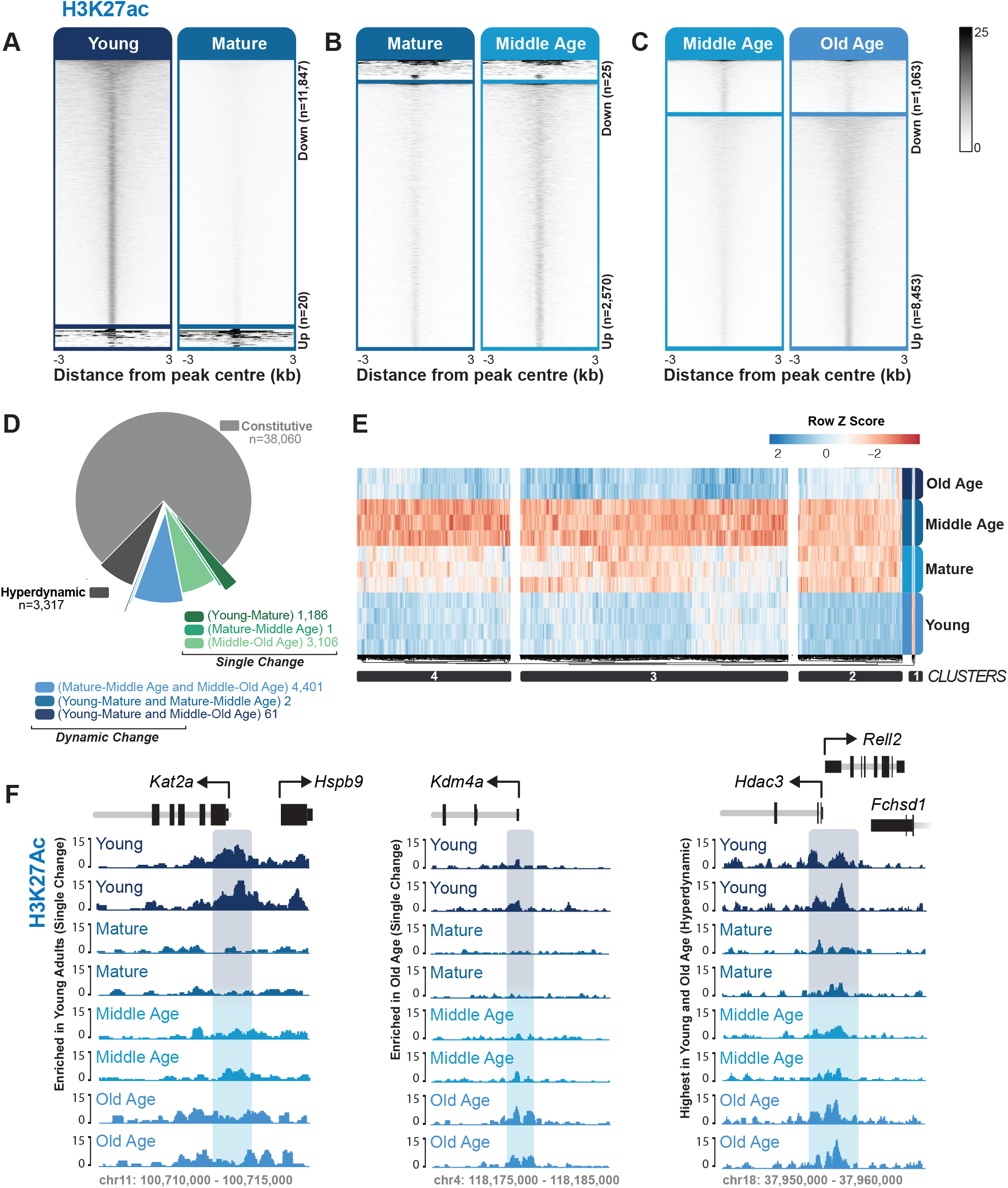
H3K27ac at Promoters. **A-C)** Heatmaps of H3K27ac signal at differential regions for (**A**) young versus mature adults, (**B**) mature versus middle age adults and (**C**) middle versus old age adults. Signal is positioned on the centre +/- 3 kb of each differential region and sorted in descending order. **D)** Percentage of promoters in each category (analysis as per Figure 3D) determine by changed in H3K27ac signal. Single and dynamic changes are further classified based on which time point the change occurred. **E)** Heatmap of normalised (row Z score) ChIP-seq signal of H3K27ac at hyperdynamic promoters. Promoters are divided into four groups by hierarchical clustering. **F)** Example regions of H3K27ac signal at epigenetic genes *Kat2a*, *Kdm4a* and *Hdac3*, displaying two representative tracks for each age.

We again categorised promoters into the four groups of change (constitutive, single change, dynamic, hyperdynamic), this time utilizing the H3K27ac enrichment signal. The majority of promoters (38,069; 76%) had no significant change in H3K27ac (Figure 5D). Of the promoters with a change in H3K27ac, there were 4,293 that exhibited a single change and interestingly, these changes occurred mostly either between young and mature adults (1,186) or between middle and old age (3,106; Figure 5D). A similar number of promoters were dynamic (4,464), with the majority also changing early (young vs mature adult neurons) and late (middle to old age) during aging (98%; Figure 5D). There were 3,317 promoters hyperdynamic for H3K27ac in neurons, which clustered into four groups by patterning (Figure 5E). Surprisingly, clusters 2-4, which encompass most of these regions, had a triphasic pattern of H3K27ac enrichment that was high in young adults and in old age and was reduced in mature adult and middle aged neurons. Each cluster varied slightly in the degree of enrichment, while following this triphasic pattern. Together, this shows that while there is a general loss of H3K27ac after young adulthood, this mark is recapitulated in old age at a subset of gene promoters.

We next performed GO analysis of the promoter groups for H3K27ac. Notably, the significant GO terms associated with a change in H3K27ac were mostly associated with a change occurring between young vs mature adults and/or middle vs old age adults (including single changes at either of these time points or dynamic changes occurring at both), as well as with hyperdynamic promoters (Extended Data Figure 6A-B, 7A). As with H3K4me3, significant GO terms related to processes that are cellular hallmarks of aging. This included DNA repair (dynamic promoters), which recapitulated the H3K27ac signal from young neurons in old age, as well as several pathways relating to mitochondrial function (Extended Data Figure 6A-B, 7A). Several significant GO terms were also related to epigenetics and chromatin regulation including chromatin organisation, RNA processing, chromatin modifications, nucleotide synthesis, histone acetylation, PcG protein complex, and SWI/SNF superfamily-type complex (Extended Data Figure 6A-B, 7A). Positive gene hits within these terms include several epigenetic histone modifiers such as a histone acetyltransferase (*Kat2a,* Figure 5F), histone deacetylases (*Hdac10, Hdac3*; Figure 5F), and lysine demethylases (*Kdm-3a, 4a, 5a, 6a* and *8*; Figure 5F). It also included RNA modifier *Nsun2,* a RNA cytosine methyltransferase (Extended Data Figure 7B), and several ATP-dependent chromatin remodelling enzymes such as from the CHD family (*Chd2, Chd5, Chd6*, *Chd8*), ISWI family (*Smarca5)* and the SWI/SNF family (*Smarca2, Smarcc2* and *Smarcd3;* Extended Data Figure 7B). The hyperdynamic H3K27ac promoters were enriched for biological processes such as modulation of synaptic transmission, cognition, learning and memory, along with cellular components relating to neuronal function including synapses, synaptic substructures and axons (Extended Data Figure 6A-B, 7A). This indicates that H3K27ac has a role in synaptic plasticity, learning and memory across lifespan. Together, this reveals biologically relevant gene networks accompanying age-related changes in H3K27ac. Notably, this included a consistent theme of epigenome-associated changes that were highly enriched in young neurons were subsequently recapitulated in old age.

### H3K27ac enriched enhancers and super-enhancers exhibit a recapitulation of early adulthood signatures in aged neurons

As H3K27ac is also enriched at enhancers, gene regulatory regions that are distal to promoters, we examined the behaviour of H3K27ac at regions distal to gene promoters. We classified distal regulatory region into three categories; single change, dynamic or hyperdynamic. We did not identify distal constitutive regions as there is no defined complete list of putative enhancers available for neurons in the aging brain. We found 4,923 distal regions that changed once across the neuron lifespan, with almost all changing either early in adult life (72% between young and mature adults) or late in life (27% between middle and old age; Figure 6A). There were 1,753 dynamic distal regions, where 99% (1,739) changed early (between young and mature adults) and late in life (between middle and old age; Figure 6A). There were also 646 hyperdynamic distal regions (Figure 6A). Of these, 96% (621 regions) had a gradual decrease in H3K27ac signal from young adults through to middle age, which then increased in old age (Figure 6B). Therefore, the single-change, dynamic and hyperdynamic distal regions also largely demonstrate the triphasic recapitulation of early adulthood in aged neurons (Figure 6C).

**Figure 6.**
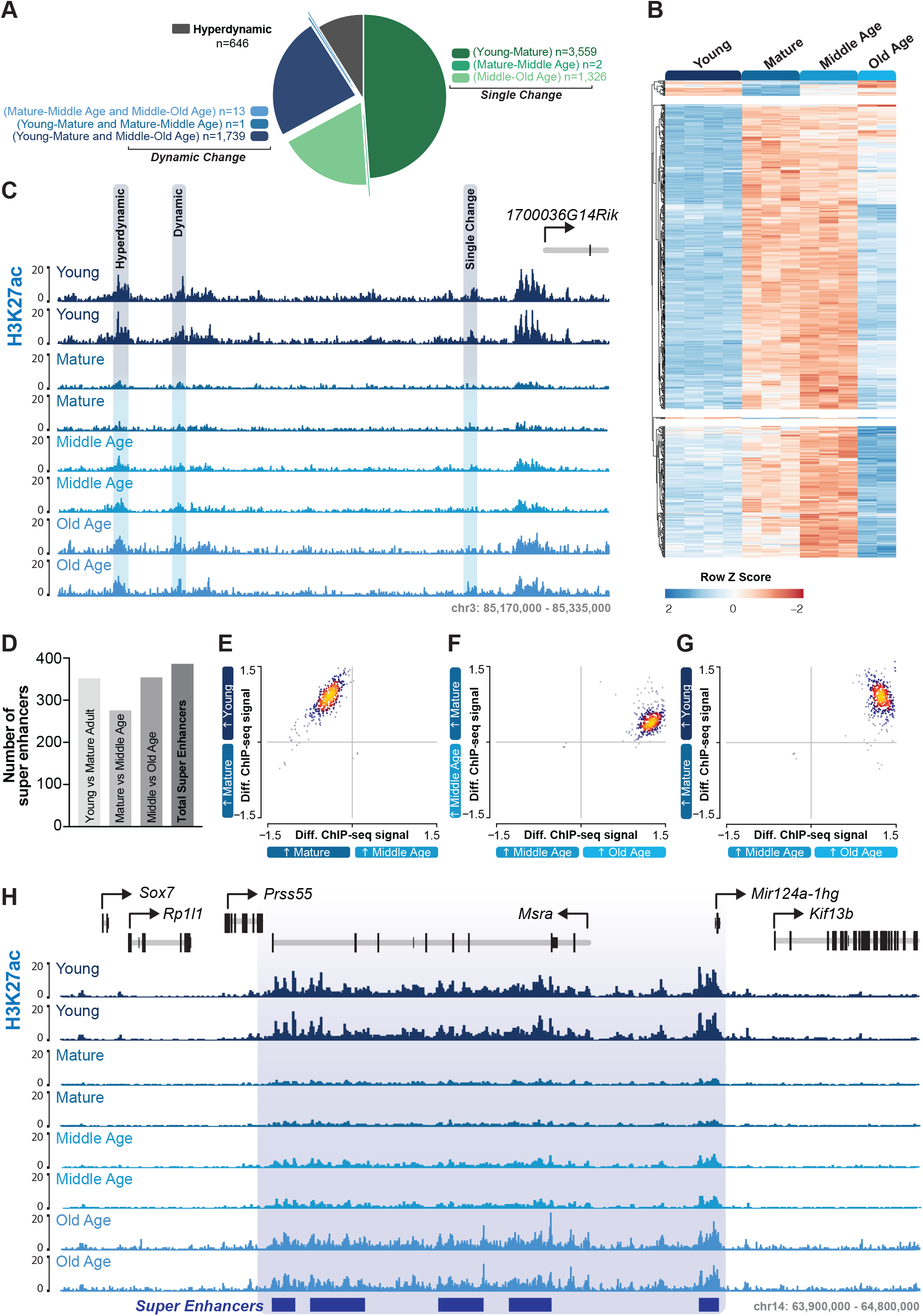
H3K27ac at enhancers and super-enhancers. **A)** Percentage of distal regions classified based on the number of times there is a change in H3K27ac signal across the mouse lifespan. **B)** Heatmap of normalised (row Z score) ChIP-seq signal of H3K27ac at hyperdynamic distal regions. Regions are divided into three groups by hierarchical clustering. **C)** Example genome browser regions of H3K27ac signal on chromosome 3 showing examples of a single change between young and mature neurons, a dynamic region which changes between young and mature adult and then again between middle and old age, and a hyperdynamic region that exhibited decreased H3K27ac marking between young and mature adult neurons, increased in middle age and then further increased in signal in old age. Two representative tracks for each age are displayed. **D)** Overlap of differentially enriched distal H3K27ac regions with neuronal super enhancers. **E)** Differential signal of H3K27ac at super enhancers for young versus mature adults (y-axis) and mature versus middle age adults (x-axis) with most of the super enhancers showing a stronger signal in the young compared to mature and middle age neurons. **F)** Differential signal of H3K27ac at super enhancers for mature versus middle age adults (y-axis) and middle age versus old age adults (x-axis) with most of the super enhancers showing a stronger signal in the old age compared to mature and middle age neurons. **G)** Differential signal of H3K27ac at super enhancers for young versus mature adults (y-axis) and middle age versus old age adults (x-axis) with most of the super enhancers showing a stronger signal in the young and old age adults compared to mature and middle age neurons. **H)** Example genome browser regions of H3K27ac signal at super enhancers within the Msra gene, showing two example tracks for each age.

Super enhancers are defined as multiple enhancers in close proximity that collectively drive gene transcription linked to cell identity. We overlapped the differential H3K27ac distal regions with annotated putative neuronal super enhancers^25^ to determine if their epigenetic signature is altered with neuronal aging. Of the 389 neuronal super enhancers, greater that 70% had altered levels of H3K27ac across the adult neuron lifespan (Figure 6D). We compared the difference in H3K27ac signal at neuronal super enhancers pairwise in a chronological fashion across our time course. We found that the H3K27ac signal at super-enhancers decreased from the young adults through to middle age, followed by an increased in old age (Figure 6E-G), such as within and around the gene *Msra* (Figure 6H). The vast majority of H3K27ac marked dynamic and hyperdynamic enhancers and super enhancers demonstrated a recapitulation of early adult enrichment in old age, suggesting an age-related program of H3K27ac plasticity potentially related to cell identity and function.

## DISCUSSION

We assembled comprehensive maps of H3K4me3 and H3K27ac epigenetic modifications in neurons across an aging time course. The most dramatic change oberseverd in our aging time course was a loss of H3K4me3 and H3K27ac marking between young adult and mature adult neurons. Neurons between 3 and 6 months of age represent the transition from young to mature adult mice ^26,27^. H3K27ac and H3K4me3 marking are generally indicative of increased transcriptional activity ^28,29^, and enrichment of these marks at gene promoters in young adult mice may reflect a critical time of neuronal plasticity and maturation^30^. The current dogma of brain maturation suggests that neuronal connections are established in mice by 3 months of age^27^, however, emerging evidence shows ongoing developmental changes beyond 3 months^26,27,30^. Our data indicate that the maturation of neurons continues between young adult and mature adult mice, with genes involved in cell proliferation and brain development losing H3K4me3 marking between 3 and 6 months of age. This is supported by reports of widespread synaptic pruning in pyramidal neurons in the cortex of mice at 3 months of age^30^, and the stabilisation of adult levels of neurotransmitters and synaptic densities after 3 months of age in mice^26^. Furthermore, the emergence of adult behavioural tendencies, such as reduced risk taking and increased parenting, occurs after 3 months of age^26,27^. Thus, enrichment of H3K27ac and H3K4me3 marking in young adult neurons may reflect the developmental plasticity critical for establishing gene expression profiles for stable neuronal networks in mature adult mice^26^.

The vast majority of promoters with plastic age-related H3K4me3 marking in neurons were hyperdynamic. Each of the 4 main hyperdynamic clusters of H3K4me3 marked promoters in neurons had peak H3K4me3 marking at a different stage of life. In contrast, promoters with changing H3K27ac marking were evenly split between the single change, dynamic and hyperdynamic categories (∼30% each). GO analysis of hyperdynamic H3K4me3 promoters and plastic H3K27ac promoters (including single change, dynamic and hyperdynamic promoters) revealed known traits of cellular and neuronal aging including DNA repair, protein folding, mitochondrial function, non-coding RNA regulation and synaptic function. This is supported by previous reports of age-related changes in H3K4me3 peaks in human neurons in synaptic structure/transmission and DNA repair pathways^20^ as well as age-related transcriptomic changes in mouse neurons that mapped to non-coding RNA processing and metabolism and mitochondrial function pathways^31^.

DNA repair pathways play a vital role in neurons, which are highly metabolically active, post-mitotic and long-lived. In aging neurons, DNA breaks and single nucleotide variants increase^24,32,33^. In this study, pathways related to DNA repair were detected in promoters with hyperdynamic H3K4me3 and dynamic H3K27ac marking, including DNA repair, base excision repair, inter-strand crosslink repair and DNA mismatch repair. We also observed H3K4me3 enrichment at the promoters of genes involved in DNA repair to decrease with age, including genes involved in non-homologous end joining repair (*Lig4, Xrcc6*) and break-induce DNA replication (*Pold3, Rad52*, *Brca2, Rad51*). Interestingly, neuronal activity induces double strand DNA breaks near early response gene promoters that is required for their expression^34,35^, and the base excision repair pathway also plays a critical role in the expression of late response genes following neuronal activity^36^. Thus, neurons implement selective DNA repair, with actively transcribed promoters being repaired more frequently^24^. Furthermore, recent data determined that DNA single-strand breaks in iPSC derived neurons were co-localised with enhancers, including 90% of super-enhancers associated with neuronal identity^37^. Thus, the changes in DNA repair related pathways in our data may be inextricably linked with the changes in transcription-independent genome maintenance as well as the changes in neuronal activity and synaptic plasticity that occur in aged neurons.

Our data also highlight that aged neurons recapitulate the H3K27ac enrichment at promoters, enhancers and super enhancers observed in young adult neurons. This mirroring of epigenetic signatures at the extremes of our aging time course mapped to genes involved in chromatin organisation/modification, protein folding/modifications, RNA processing, and synaptic function and structure. Synapse loss and alterations in synaptic function and plasticity occur with age^1,38^. Thus, it is possible that pathways related to synaptic plasticity are reactivated in aged neurons with the aim of maintaining neuron function^39,40^ and combatting age-associated cognitive decline. The concept of age associated recapitulation of a youthful epigenetic state has been proposed in previous literature^41,42^. Studies have shown that the epigenome is divergent throughout life ^10^, but cortical aging leads to a convergence of inter-individual DNA methylation states, suggesting cell de-differentiation in the aging human frontal cortex^41^. Of note, enhancers and super enhancers contribute to cell type specificity, particularly in the brain^43^, and the reversion of enhancers and super enhancers to a young adult status may contribute to cell de-differentiation. While the role of developmental programs in aging is contentious, there is increasing evidence that developmental programs may play an important role in aging alongside the accumulation of molecular damage over time^44–46^. Transcriptomic data from the human prefrontal cortex indicates that miRNA and transcription factors regulate gene expression programs, not just during development, but across lifespan; inextricably linking development and aging^45,46^.

Chromatin organisation and modification were a common theme identified with H3K27ac recapitulation in neuronal aging. The promoter of histone acetyl transferase *Kat2a* exhibited altered H2K27ac enrichment between young and mature adult neurons. *Kat2a* has critical activity in transcription, where it targets multiple sites on histone-3 for acetylation^47^. The reduction of H3K27ac at the promoter of *Kat2a* is suggestive of a positive feedback loop, where down regulation of *Kat2a* is concomitant with a reduction in H3K27ac genome wide. Some studies have reported that *Kat2a* inhibition promotes cellular lifespan^47,48^, and others have linked *Kat2a* reduction to impaired hippocampal synaptic plasticity and long-term memory consolidation^49^. Interestingly, we also observed changes in H3K27ac at the promoters of histone deacetylases *Hdac3* and *Hdac10*. The overexpression of *Hdac3* in the rat brain causes death of otherwise healthy neurons and it is associated with preventing long-term memory formation^4,50^. Concomitantly, reduced expression protects against neuronal death^50^, and rescues long-term memory impairments^5^. The targets of *Hdac10* have not been well defined, however, it has been associated with the activity of *Hdac3*^51,52^. We also identified several ATP-dependent chromatin remodelling complex subunits, in particular members of the SWI/SNF complex and CHD family, in chromatin organisation pathways enriched in young and old aged neurons. Both SWI/SNF and CHD remodellers have been associated with neuronal development^53–55^, for example, *Chd5* is specific to the brain and mice lacking *Chd5* have autism-like characteristics^53^. However, there is little research into the role of ATP-dependent remodellers in the aging brain. *Chd5* has been associated with the regulation of neuronal genes linked to aging and Alzheimer’s disease^56^, but the effect of *Chd5* on the epigenetic signature of these genes during the normal aging process is unclear. This highlights a link in Chd5 function and epigenome regulation between development and aging in the brain.

Our study revealed decreasing rDNA chromatin regulation as a characteristic of neuron aging (Extended data Figure 2). rDNA encodes the proteins necessary to produce ribosomes and plays roles in genome stability, widespread epigenetic regulation of transcription and lifespan^57,58^. In mice, there are hundreds of copies of rDNA in tandem repeat arrays, a subset of which are required for the production of ribosomes, and the remainder is silenced in heterochromatin^59^. rDNA loci are inherently unstable, and in dividing cells copy loss naturally occurs during aging and is combated using homologous recombination^59,60^. In the aging mouse brain and human cerebral cortex no change of rDNA dose or copy number have been detected^61,62^. To date, there have been no studies of rDNA in aging neurons, however, our data highlights that rDNA regulation warrants further investigation specifically in aging neurons.

In summary, to address a need for neuronal specific epigenetic information, our study has generated reference maps of H3K4me3 and H3K27ac from neuronal nuclei across the adult mouse lifespan. This catalogue of active promoters and putative enhancers demonstrate the plasticity of the epigenome in aging and identified rDNA and epigenetic regulation as novel hallmarks of neuron aging.

## METHODS

### Mouse tissue processing

All animal procedures were undertaken with ethical approval from the Animal Ethics Committee of the University of Tasmania (A12780/A15120) and all experiments abided by the Australian Code of Practice for the Care and Use of Animals for Scientific Purposes. Male mice aged 8-10 months were used for ChIP-qPCR analysis of whole brain (n=3) and isolated neuronal nuclei (n=2). Male mice for ChIP-seq were aged to 3 months (young adult), 6 months (mature adult), and 12 months (middle age adult) and 24 months (old age adult), with n = 5 mice per time point. Animals were housed in standard conditions (12hr day/night cycle, housing temperature of 20°C, *ad libitum* access to food). Mice were sacrificed with an intraperitoneal injection of Lethobarb (110mg/kg), and cardiac perfusion with 0.01M PBS. The forebrain was then rapidly dissected and snap frozen in liquid nitrogen.

### Isolation of nuclei from mouse forebrain

Fresh-frozen mouse forebrains were sectioned along the midline. Half of each forebrain was immersed in 4.8 ml nuclei extraction buffer (NEB [Sucrose 0.32M, CaCl 5mM, Mg(Ac).4H_2_0 3mM, EDTA 0.1mM, Tris-HCl pH8.0 10mM, 1x protease inhibitor cocktail, PMSF 0.1mM, Triton-X-100 0.1%]) on ice and homogenised with a dounce tissue homogeniser (#357544, Wheaton, Millville, New Jersey, USA). Homogenised tissue was filtered through four layers of cheesecloth (#9338918004577, Ogilvies Designs, Belmont, Western Australia, Australia), followed by 70μm and 40μm cell strainers (#352350, #352340, Thermo Fisher Scientific, Waltham, MA, USA) on ice. Nuclei were pelleted by centrifugation and resuspended 500 μl 0.01M PBS. Nuclei (90%) were incubated with anti-NeuN antibody (1:1000, #MAB377, Merk Millipore, Billerica, Massachusetts, USA) and goat-anti-mouse AlexaFluor 647 (1:2000, #A31571, Invitrogen/Life Technologies, Carlsbad, CA, USA) in blocking solution (0.5% BSA, #B4287 25G, Sigma-Aldrich, St Louis, Missouri, USA; and 10% normal goat serum, #G9023-10, Sigma81 Aldrich, St Louis, Missouri, USA in 0.01M PBS) for 40 mins at 4°C on a slow rotator in the dark. The remaining 10% of the nuclei were used as the secondary only control and were incubated with goat-anti-mouse AlexaFluor 647 (1:2000) in blocking solution for 40 mins at 4°C on a slow rotator in the dark. DAPI (#D3571, Life Technologies, Carlsbad, CA, USA) was added to the secondary only control for the final 5 mins of the incubation. Following antibody incubations, nuclei were collected by centrifugation then resuspended in ice cold PBS.

### Fluorescence activated nuclei sorting (FANS) for neuronal nuclei

FANS was performed on a BD Biosciences FACS Aria III (Becton Dickinson Biosciences, Franklin Lakes, NJ, USA). The secondary antibody only negative controls established gating for each sample and NeuN labelled nuclei were then sorted and collected in tubes pre-coated with foetal calf serum (FCS; #SH30084.02, Hyclone, GE Healthcare Life Sciences, South Logan, Utah, USA). Purity checks were performed and all samples were validated at >97% purity.

### Chromatin immunoprecipitation (ChIP)

NeuN+ nuclei were fixed in 1% methanol-free formaldehyde (#28906, Pierce, Thermo Fisher Scientific, Waltham, MA, USA) for 15 mins at room temperature then quenched with 0.125M glycine (#G8898, Sigma-Aldrich, Missouri, USA) for 5 mins, prior to freezing at −80°C. Neuronal nuclei were the collected by centrifugation at 52,000 × g at 4°C for 25 mins (Sorvall WX Ultra 90 ulracentrifuge; TH-641 swinging bucket rotor (#54295, Thermo Fisher Scientific, Waltham, MA, USA), resuspended in SDS lysis buffer (EDTA 10mM, SDS 1%, Tris-HCl pH8.1 50mM) and transferred to 1.5mL Bioruptor+ TPX microtubes (#C30010010-300, Diagenode, Seraing, Belgium). Samples were sonicated on the Bioruptor Plus next-gen ultrasonicator (Diagenode, Seraing, Belgium) for 50 cycles of 30 secs on / 30 secs off, on ice. Chromatin fragment size was confirmed to be 200-500bp on the Agilent 4200 tape-station with HS D1000 tape and reagents (#5067-5584, Agilent Technologies, California, USA). Chromatin immunoprecipitation was performed as previously described previously^63–65^ with minor modifications. Briefly, 5 × 10^5^ nuclei were used per immunoprecipitation. All samples (n = 20) were processed in parallel to eliminate batch effects. Samples were incubated with primary antibodies anti-H3K4me3 (2μg; #39160; Active Motif, Carlsbad, CA, USA) or anti-H3K27ac (2μg; #39134; Active Motif, Carlsbad, CA, USA) overnight at 4°C with rotation. All antibodies have been previously validated for specificity and use in ChIP assays.

### ChIP-qPCR

DNA from whole brain and neuronal nuclei ChIP samples was purified with the Chromatin IP DNA purification kit (Active Motif, #58002) according to the manufacturer’s instructions. Real-time quantitative PCR was performed using 5uL of KAPA Sybr Fast Universal 2x PCR master mix (Merck, #KK4601), 0.6uL of 5 uM forward primer, 0.6uL of 5uM reverse primer, 2uL sample and 1.8uL of nuclease free water per reaction. Primer sequences targeting promoter regions were as follows; *Sorbs3* For-GCCCAACAAGGAAGAGAGC, Rev-CAAGTCCCAAACTCCTCCTG, *Picalm1* For-GACGAGCCGCAGAGATGT, Rev-GTCGTGGCCTTGCATACTGT and *Bin1* For-CCCCTGAGCTGTTCTAGTGC, Rev-GAAGCGAGCGCGGATAAT. Signal was calculated as enrichment over IgG and over total input (%).

### ChIP library preparation and next-generation sequencing

Sequencing libraries were prepared using the Nugen Ovation Ultralow V2 (#0347 V2 1-96/0344 V2 1-16, Redwood City, CA, USA) library preparation kits as per the manufacturer’s protocol. Library quality control and normalisation was carried out by the Australian Genomics Research Facility with the Agilent 4200 tape-station system and KAPA qPCR quantification. Samples were then pooled and distributed equally across all sequencing lanes. Samples were sequenced on the Illumina Hi-Seq 2500 Next-Generation-Sequencer (Illumina, San Diego, CA, USA) using 50bp single-end sequencing.

### ChIP-seq sequence alignment and pseudoreplicate generation

Sequencing FASTQ files were imported onto the Nectar Research Cloud for all processing. Raw reads were checked for quality control with FASTQC version 0.11.7 and MultiQC^66,67^. Adapters were trimmed using the Trim Galore wrapper version 0.4.3 for Cutadapt (version 1.15)^68^. Samples were aligned with Bowtie2^69^ version 2.3.4 to *Mus musculus* reference genome mm10 using default parameters^70^. After mapping, samples were converted to BAM file format and indexed with SAMtools version 1.7. Bigwig files were generated with bamCompare from deepTools for visualisation on IGV^71,72^. Samples not passing QC were removed at this stage, with between 2-5 samples retained for each histone mark at each time point. Reads from samples for each mark at each time point were pooled using samtools *merge,* read with samtools *view*, shuffled, and randomly distributed into pseudoreplicates equal to the number of samples passing QC to obtain an average signal for the mouse population assayed^73,74^.

### PCA and Pearson Correlation of Pseudoreplicates

The log counts per million ChIP-seq signal for each sample was determined for 500bp bins across the entire mm10 genome using *Subread*. This data was then used to perform PCA analysis using ‘prcomp’ from the *stats* package in R. and the first and second dimensions were plotted. The Pearson correlation coefficients were then calculated between each sample and plotted as a matrix using *ggplots2*.

### Peak calling and genome coverage

Peaks were called for H3K4me3 and H3K27ac against a merged input control utilising MACS2 with a Q value cut-off of 0.05^75^. Any ENCODE blacklisted regions^76^ for mm10 were removed from the called peaks. Percent of genome covered was calculated by accumulating the number of base pairs covered by all called peaks within a sample, divided by the total number of base pairs within the mm10 genome and multiplied by 100.

### Differential analysis

Differential enrichment analysis was performed within R using the ChIP-seq Analysis with Windows (csaw) package^77^. Briefly, bam files from pseudoreplicates were imported with the following parameters: *frag.length: 250, window.width 150, spacing 50, minq =30*, then filtered to remove blacklisted regions^76^. Samples were counted into 2000bp bins for further filtering of low abundance regions of non-specific binding, keeping reads higher than log_2_(3) above background. After filtering steps, reads were binned into 10kb bins for removal of library composition and trended bias with the scaling normalisation strategy of trimmed mean of M values (TMM) from *EdgeR*^77–79^. Differential enrichment analysis was then performed with a quasi-likelihood framework and negative binomial modelling from the *EdgeR* package^78^. Binned reads were then clustered into 2000 bp tolerance regions for H3K4me3 and H3K27ac, and false discovery rate error control was established with Benjamini Hochberg correction for multiple testing (F<0.05). Heatmaps of differential regions were created using *computeMatrix* and *plotHeatmap* tools from deepTools^80^. Heatmaps were plotted as +/- 3 kb from the centre of each differential region and sorted in descending order.

### ChIP-seq overlap with annotated gene regulatory features

Using the *AnnotationHub* package in R, we obtained a list of Ensembl annotated TSS in the mm10 genome. We classified the proximal promoter of these TSS by obtaining the region 1,500 bp upstream and 500 bp downstream. Differential regions for each histone mark were merged into a single consensus set using *countOverlaps*. The consensus differential regions were intersected with the annotated promoters using *subsetByOverlaps* and classified by the number of times a significant change was detected at each promoter. Heatmaps of the hyperdynamic groups were generated by calculating the number of reads per promoter using *Subread* and plotting the normalised Z-score with hierarchical clustering using *pheatmap*. clusters were extracted using *cutree* and exported for gene ontology analysis.

### Gene Ontology

Differential regions that overlapped with promoters were analyzed for enrichment of gene ontology terms in GREAT ^81^, using the whole genome as background and assigned to the single nearest gene. All significant GO terms for Molecular Function, Biological Process or Cellular Component and mouse phenotype gene sets are reported.

### Super enhancers

We obtained a list of mouse neuron super enhancers from DB super^25^. All differential distal H3K27ac regions were overlapped with the super enhancers using *subsetByOverlaps*. The ChIP-seq signal within each super enhancer was determined by the read counts for H3K27ac calculated with *Subread*. The differential ChIP-seq signal was then plotted using *heatscatter*.

### Data availability

The data generated in this study will be made available on GEO. The neuronal super enhancer regions are available from DB super^25^.

## ACKNOWLEDGEMENTS

We thank Dr Liz Caldon, Dr Rob Solomon and the Garvan Flow Cytometry facility for the FANS infrastructure and sample processing. We thank Prof James Vickers for helpful comments on the manuscript. This work is supported by grants awarded from the NHMRC (GNT1161768), ARC (IN180100005) and the Judith Jane Mason and Harold Stannett Williams Grant from The Mason Foundation. P.C.T. was supported by a fellowship (GNT1109696) and investigator grant from the NHMRC (GNT1176417). A.W. was supported by a Postdoctoral Fellowship in Dementia from the Dementia Australia Research Foundation and a fellowship from the NHMRC (GNT0544940). K.A.G. acknowledges support from the ARC (DP210103885) and the NHMRC (1185870). A.J.P. was supported by a scholarship from the Dementia Australia Research Foundation and the Yulgilbar Foundation. M.D.R. acknowledges support from the University Research Priority Program Evolution in Action at the University of Zurich. This research was supported by use of the Nectar Research Cloud, a collaborative Australian research platform supported by the NCRIS-funded Australian Research Data Commons (ARDC).

## AUTHOR CONTRIBUTIONS

Conceptualization, T.R.M, M.D.R, A.W. and P.C.T.; Methodology, A.J.P., K.A.G., A.W. and P.C.T.; Investigation, K.A.G., A.J.P., S.N.H., A.W. and P.C.T.; Visualization – K.A.G., J.M.C. and P.C.T.; Software – K.A.G.; Formal Analysis – K.A.G., J.M.C. and A.J.P; Data Curation – K.A.G. and A.J.P.; Writing – Original Draft, K.A.G and A.W.; Writing – Review & Editing, K.A.G., A.J.P., J.M.C., S.N.H., T.R.M, M.D.R., A.W. and P.C.T.; Funding Acquisition, M.D.R, T.M.R., A.W. and P.C.T.; Resources – A.W. and P.C.T.; Supervision – A.W. and P.C.T.

## COMPETING INTERESTS STATEMENT

The authors declare that they have no competing interests. The manuscript has been approved by all authors.

## EXTENDED DATA FIGURE LEGENDS

**Extended Data Figure 1.**
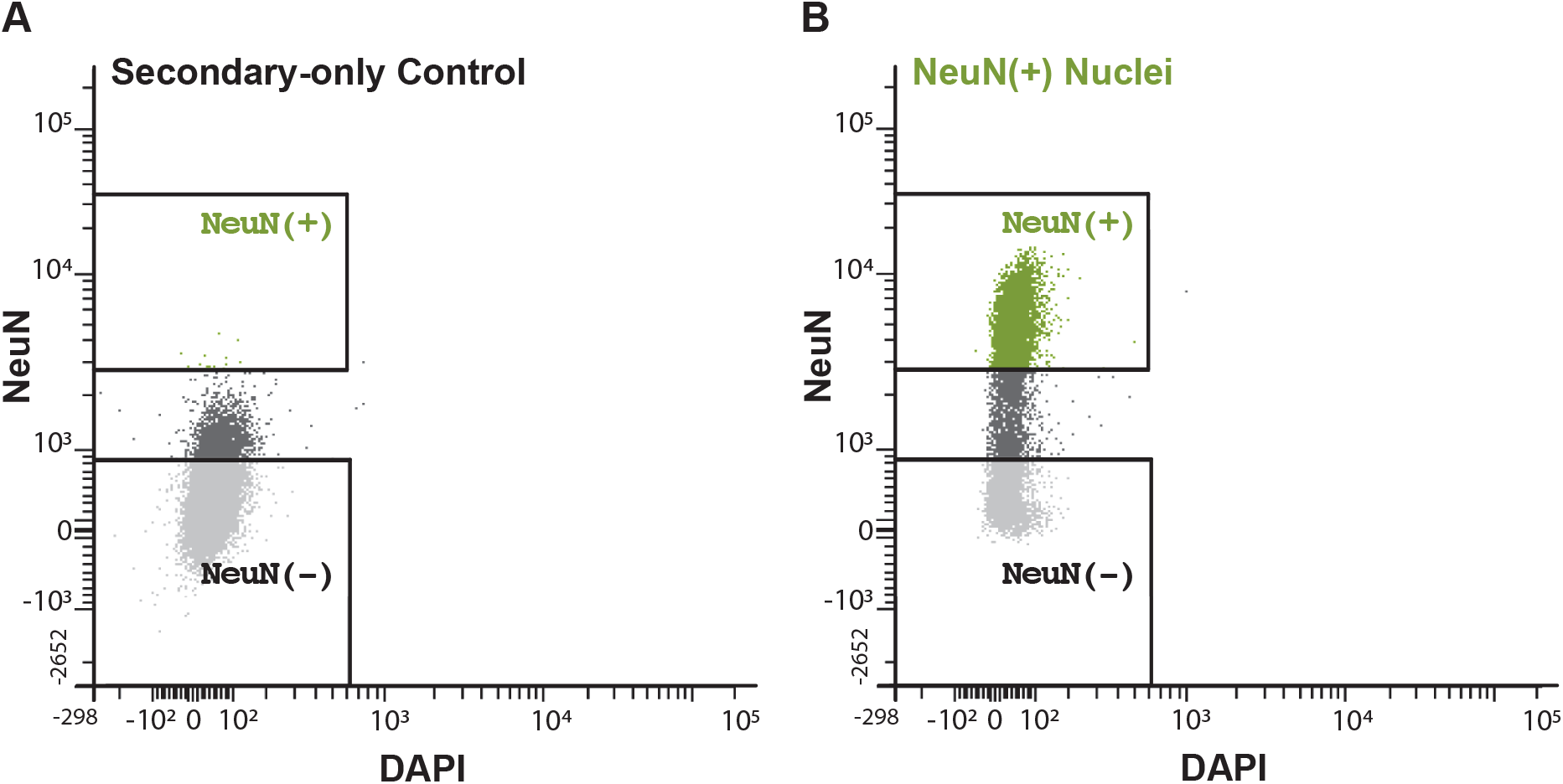
**A-B)** Gating strategy for secondary only control (A) and for purifying NeuN+ (B) mouse forebrain nuclei.

**Extended Data Figure 2.**
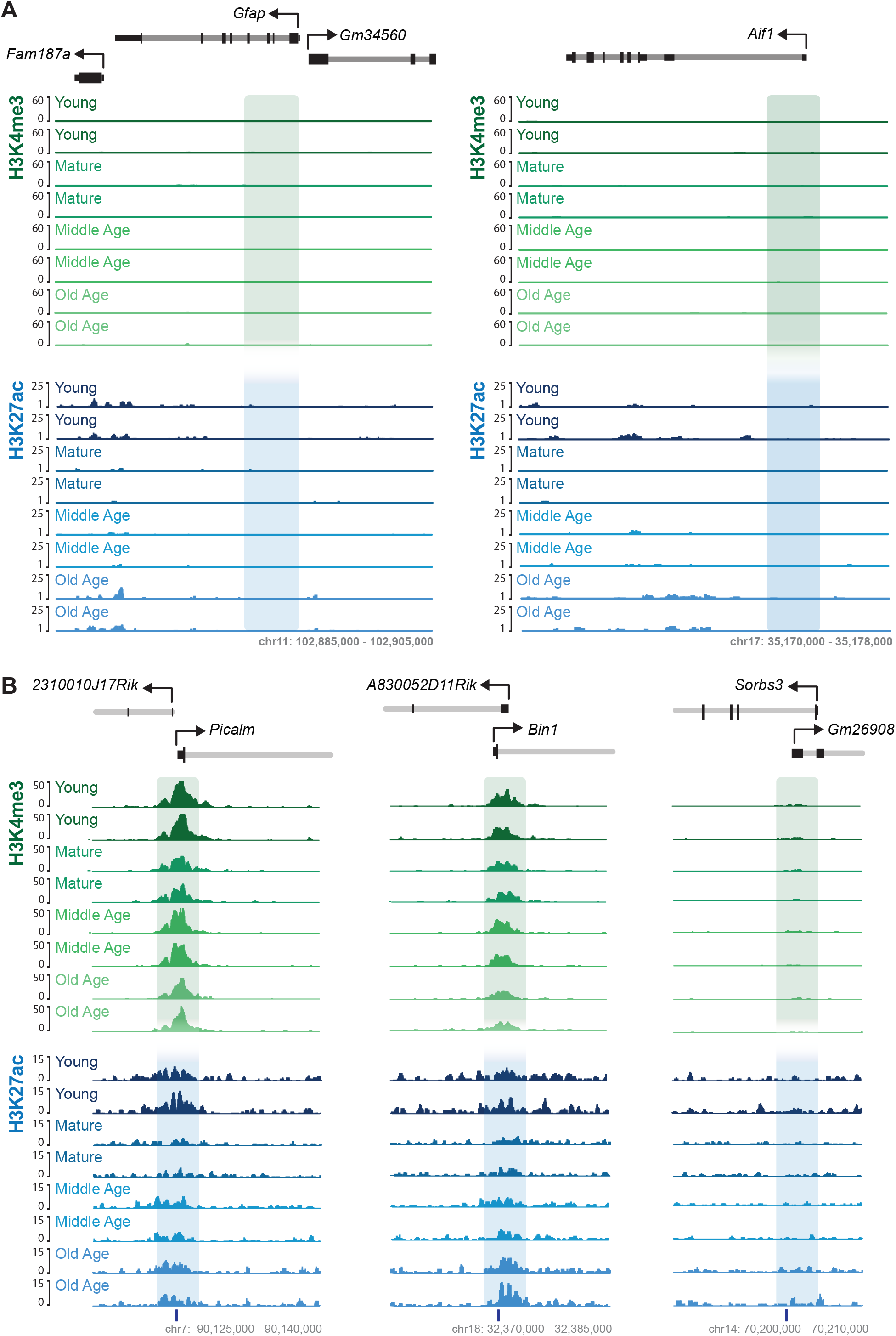
**A)** Example regions of H3K4me3 and H3K27ac neuronal ChIP-seq signal at glial specific genes *Gfap* and *Aif1,*displaying two representative tracks for each age. **B)** ChIP-seq signal of H3K4me3 and H3K27ac at the promoters of *Picalm*, *Bin1* and *Sorbs3*. Two representative tracks for each age, track underneath the ChIP-seq signal denotes the location of the primers targeting these promoters in Figure 1A-C.

**Extended Data Figure 3.**
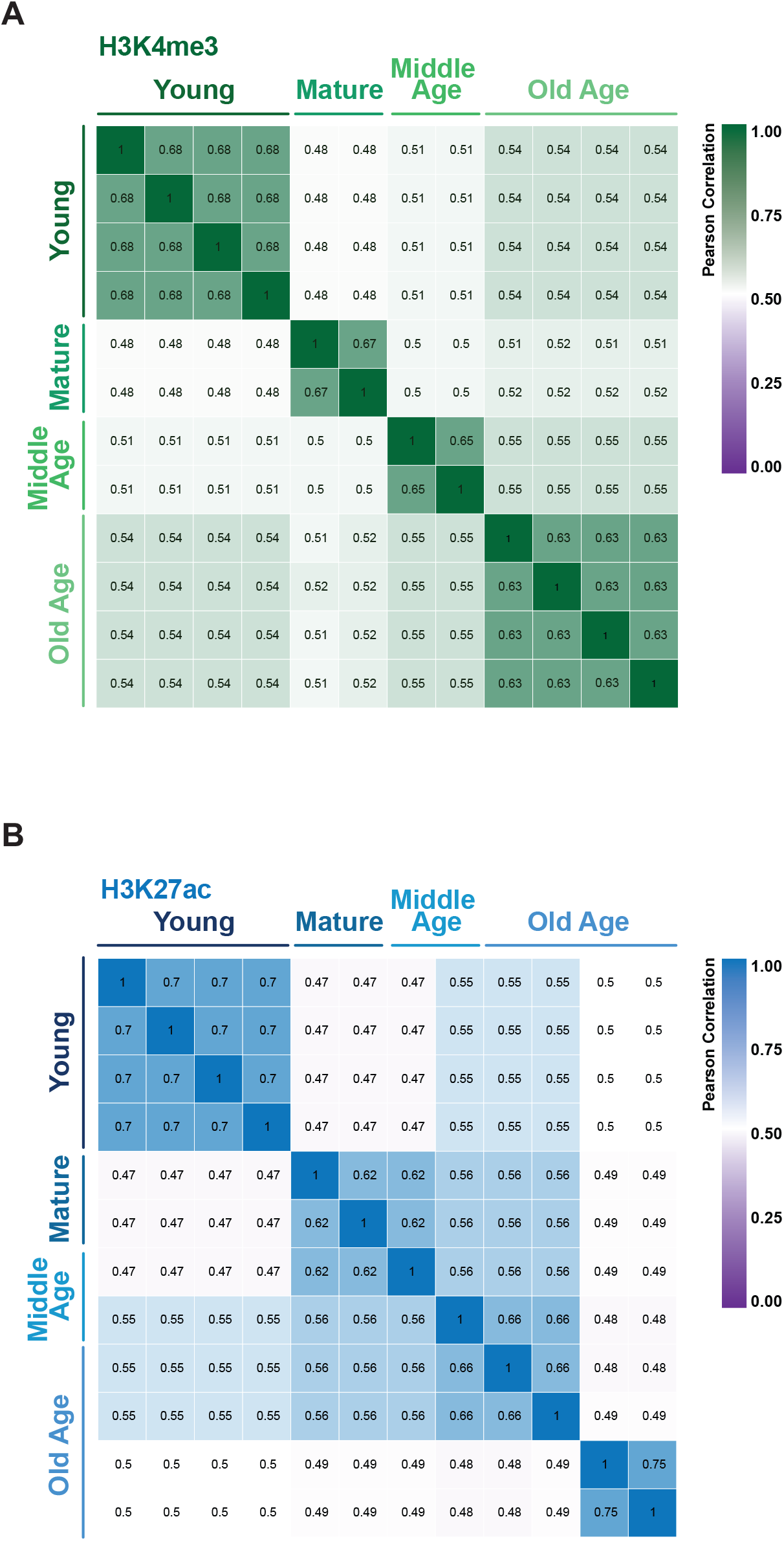
**A-B)** Matrix plots of Pearson correlations between samples for H3K4me3 (**A**) and H3K27ac (**B**). Heatmap is colour according to strength of the correlation.

**Extended Data Figure 4.**
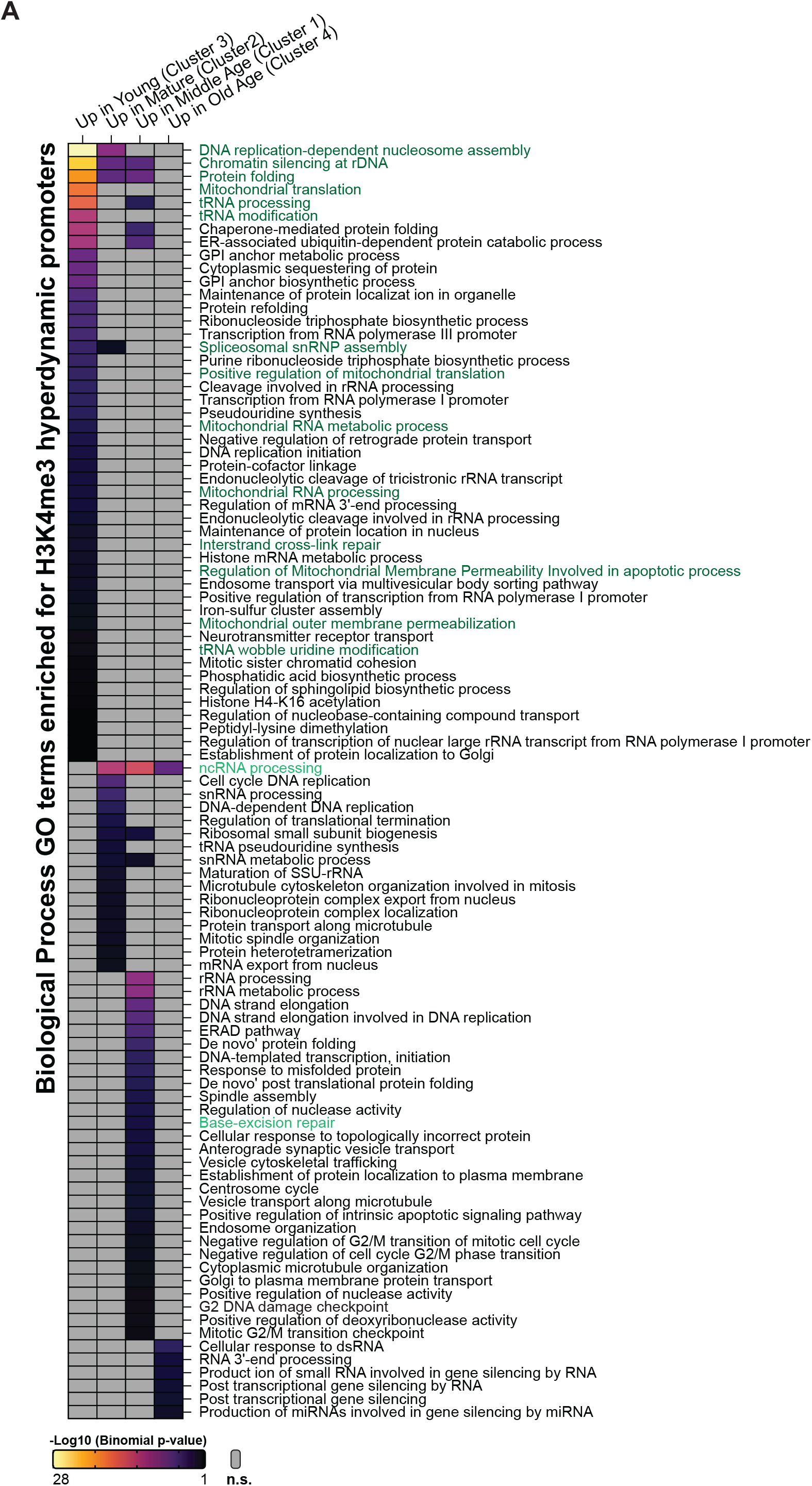
**A)** Biological process GO terms significant for the hyperdynamic promoters with a change in H3K4me3, where H3K4me3 is enriched in the young adults (Figure 4A, cluster 3), the mature adults (Figure 4A, cluster 2), the middle aged (Figure 4A, cluster 1) and old aged (Figure 4A, cluster 4) adults. Pathways discussed in the manuscript are denoted in green.

**Extended Data Figure 5.**
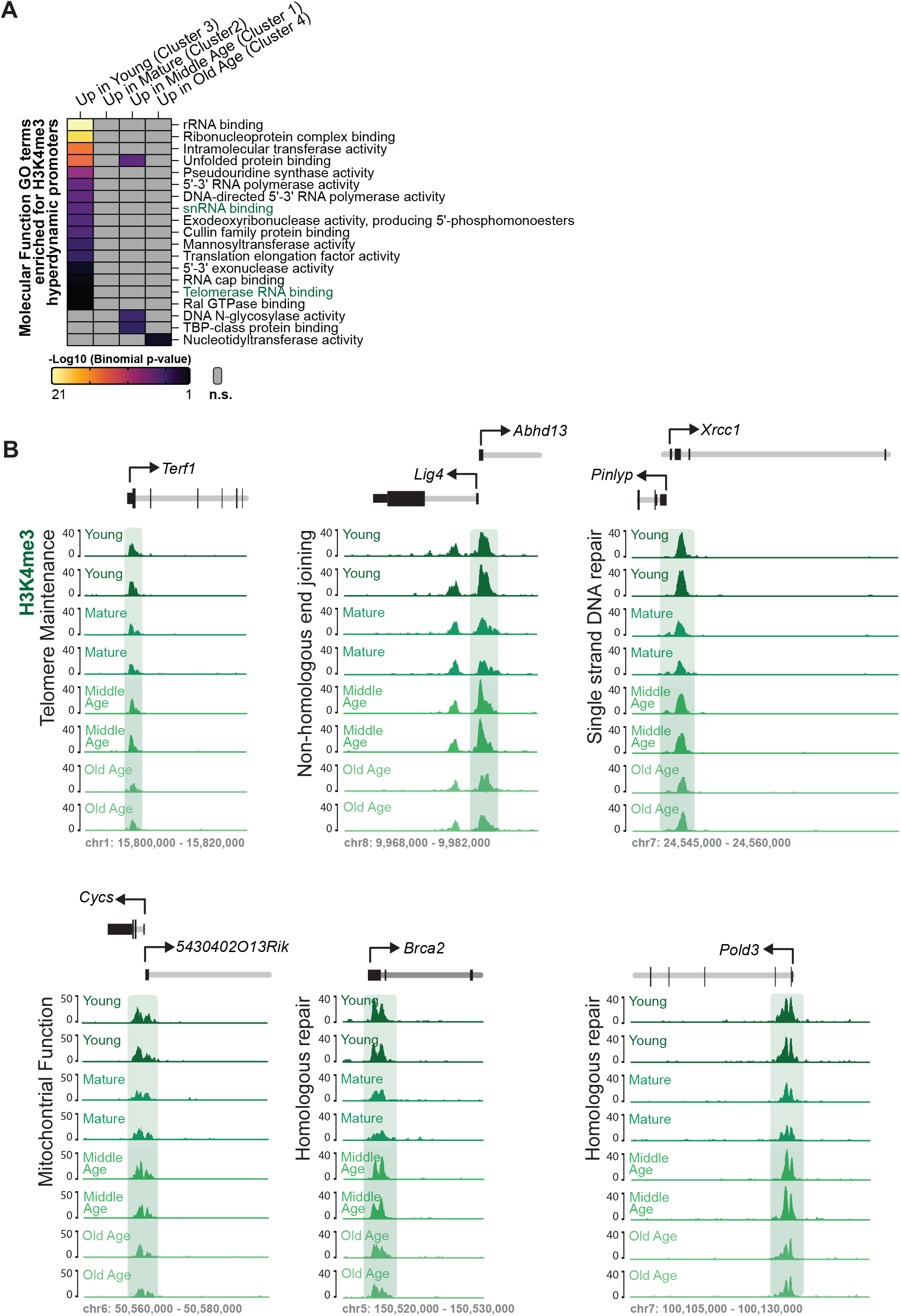
**A)** Molecular function GO terms significant for the hyperdynamic promoters with a change in H3K4me3, where H3K4me3 is enriched in the young adults (Figure 4A, cluster 3), the mature adults (Figure 4A, cluster 2), the middle aged (Figure 4A, cluster 1) and old aged (Figure 4A, cluster 4) adults. **B)** Example regions showing the H3K4me3 signal at promoters of genes involved in Telomere maintenance (*Terf1*), mitochondrial function (*Cycs*) and DNA repair (*Lig4*, *Xrcc1*, *Brca2* and *Pold3*).

**Extended Data Figure 6.**
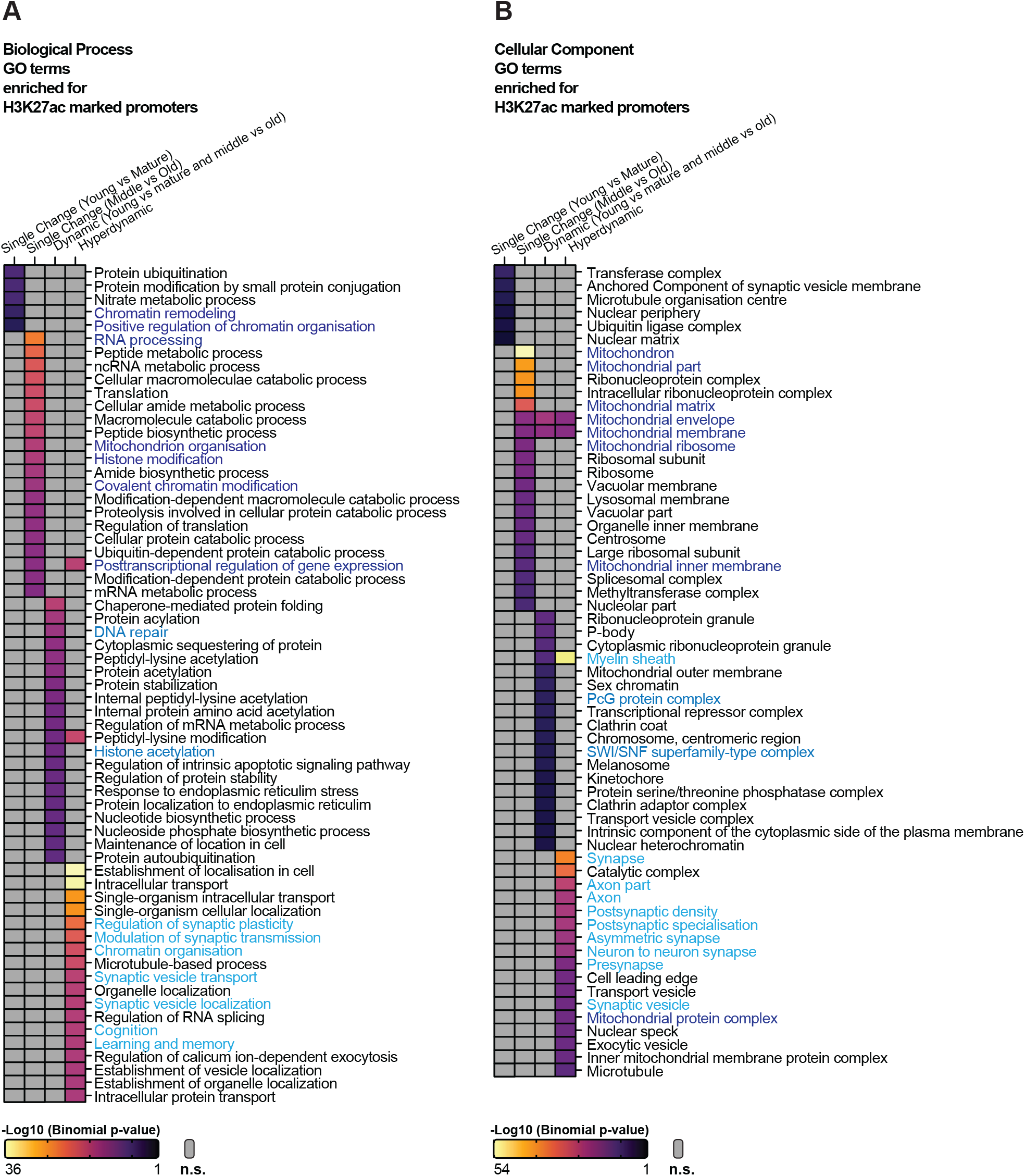
**A)** Biological process GO terms associated with a significant change in H3K27ac at promoters that has a single change between young and mature adults, a single change between middle and old aged adults, were dynamic and changed between young and mature adults as well as between middle and old aged adults, and the promoters which were hyperdynamic. Terms in blue are associated with neuronal function and epigenetic regulation. **B)** Cellular component GO terms associated with a significant change in H3K27ac at promoters that has a single change between young and mature adults, a single change between middle and old aged adults, were dynamic and changed between young and mature adults as well as between middle and old aged adults, and the promoters which were hyperdynamic. Terms in blue are associated with neuronal function, epigenetic regulation, and mitochondrial function.

**Extended Data Figure 7.**
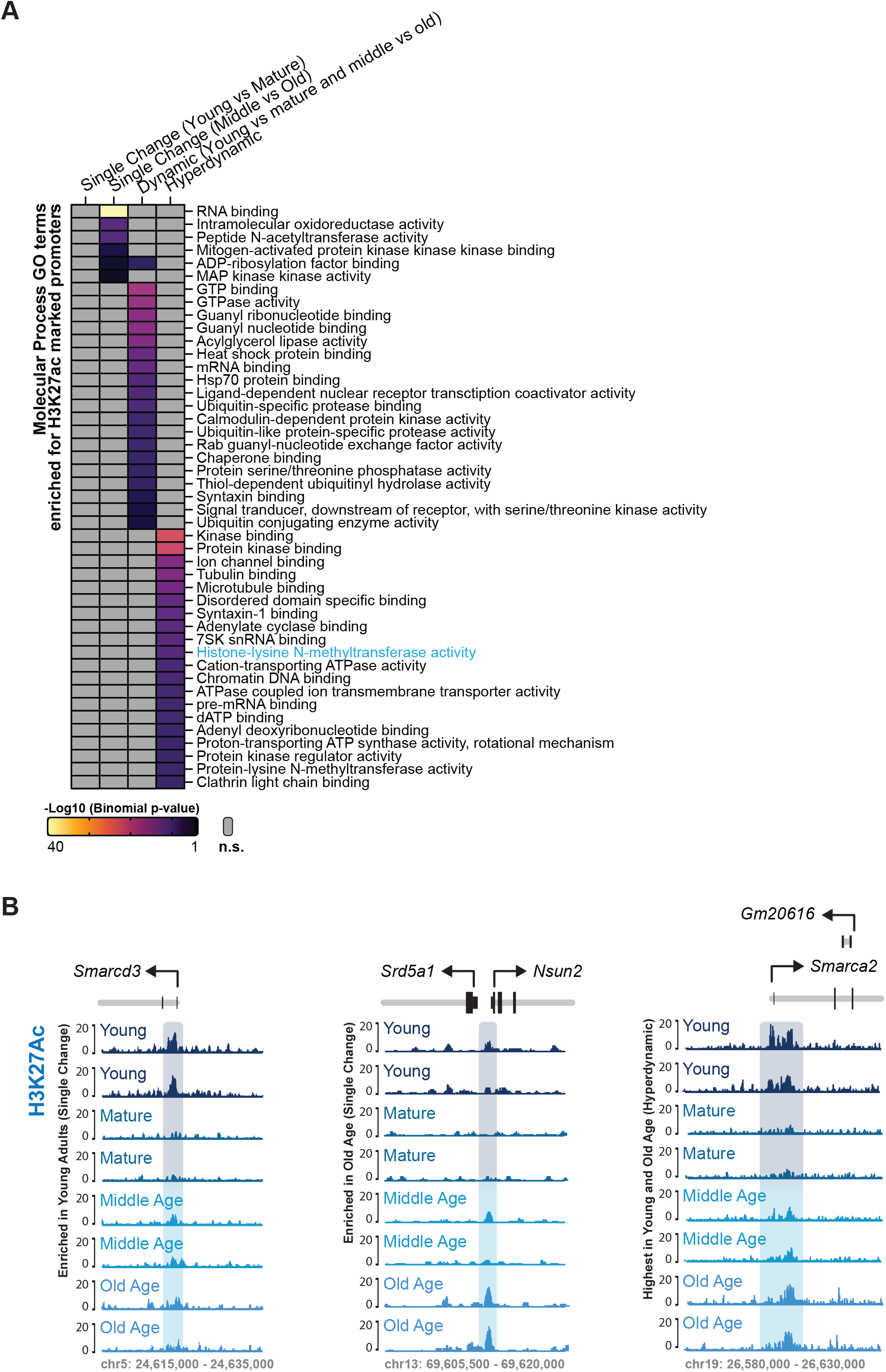
**A)** Molecular function GO terms associated with a significant change in H3K27ac at promoters that has a single change between young and mature adults, a single change between middle and old aged adults, were dynamic and changed between young and mature adults as well as between middle and old aged adults, and the promoters which were hyperdynamic. Terms associated with epigenetic regulation are shown in blue. **B)** Example regions of H3K27ac signal at epigenetic genes *Nsun2*, *Smarcd3* and *Smarca2*, displaying two representative tracks for each age.

## Notes

### Competing Interest Statement

The authors have declared no competing interest.

## REFERENCES

1 Murman, D. L. The Impact of Age on Cognition. Semin Hear 36, 111–121, doi:10.1055/s-0035-1555115 (2015).

2 Oh, M. M. & Disterhoft, J. F. Learning and aging affect neuronal excitability and learning. Neurobiol Learn Mem 167, 107133, doi:10.1016/j.nlm.2019.107133 (2020).

3 Oliveira, A. M., Hemstedt, T. J. & Bading, H. Rescue of aging-associated decline in Dnmt3a2 expression restores cognitive abilities. Nat Neurosci 15, 1111–1113, doi:10.1038/nn.3151 (2012).

4 McQuown, S. C. et al. HDAC3 is a critical negative regulator of long-term memory formation. J Neurosci 31, 764–774, doi:10.1523/JNEUROSCI.5052-10.2011 (2011).

5 Shu, G. et al. Deleting HDAC3 rescues long-term memory impairments induced by disruption of the neuron-specific chromatin remodeling subunit BAF53b. Learn Mem 25, 109–114, doi:10.1101/lm.046920.117 (2018).

6 Giles, K. A. & Taberlay, P. C. The Role of Nucleosomes in Epigenetic Gene Regulation. Clinical Epigenetics, 87–117 (2019).

7 Sen, P., Shah, P. P., Nativio, R. & Berger, S. L. Epigenetic Mechanisms of Longevity and Aging. Cell 166, 822–839, doi:10.1016/j.cell.2016.07.050 (2016).

8 Benito, E. et al. HDAC inhibitor–dependent transcriptome and memory reinstatement in cognitive decline models. The Journal of Clinical Investigation 125, 3572–3584, doi:10.1172/JCI79942 (2015).

9 Nativio, R. et al. Dysregulation of the epigenetic landscape of normal aging in Alzheimer’s disease. Nature Neuroscience, doi:10.1038/s41593-018-0101-9 (2018).

10 Cheung, I. et al. Developmental regulation and individual differences of neuronal H3K4me3 epigenomes in the prefrontal cortex. Proc Natl Acad Sci U S A 107, 8824–8829, doi:10.1073/pnas.1001702107 (2010).

11 Benayoun, B. A. et al. Remodeling of epigenome and transcriptome landscapes with aging in mice reveals widespread induction of inflammatory responses. Genome Res 29, 697–709, doi:10.1101/gr.240093.118 (2019).

12 Chouliaras, L. et al. Consistent decrease in global DNA methylation and hydroxymethylation in the hippocampus of Alzheimer’s disease patients. Neurobiology of aging 34, 2091–2099, doi:10.1016/j.neurobiolaging.2013.02.021 (2013).

13 Dagnas, M., Guillou, J. L., Prevot, T. & Mons, N. HDAC inhibition facilitates the switch between memory systems in young but not aged mice. J Neurosci 33, 1954–1963, doi:10.1523/JNEUROSCI.3453-12.2013 (2013).

14 Peleg, S. et al. Altered histone acetylation is associated with age-dependent memory impairment in mice. Science (New York, N.Y.) 328, 753–756, doi:10.1126/science.1186088 (2010).

15 Vecsey, C. G. et al. Histone deacetylase inhibitors enhance memory and synaptic plasticity via CREB:CBP-dependent transcriptional activation. J Neurosci 27, 6128–6140, doi:10.1523/JNEUROSCI.0296-07.2007 (2007).

16 Lister, R. et al. Global Epigenomic Reconfiguration During Mammalian Brain Development. Science (New York, N.Y.) 341, doi:10.1126/science.1237905 (2013).

17 Gasparoni, G. et al. DNA methylation analysis on purified neurons and glia dissects age and Alzheimer’s disease-specific changes in the human cortex. Epigenetics & Chromatin 11, 41, doi:10.1186/s13072-018-0211-3 (2018).

18 Luo, C. et al. Single-cell methylomes identify neuronal subtypes and regulatory elements in mammalian cortex. Science (New York, N.Y.) 357, 600–604, doi:10.1126/science.aan3351 (2017).

19 Mo, A. et al. Epigenomic Signatures of Neuronal Diversity in the Mammalian Brain. Neuron 86, 1369–1384, doi:10.1016/j.neuron.2015.05.018 (2015).

20 Shulha, H. P., Cheung, I., Guo, Y., Akbarian, S. & Weng, Z. Coordinated cell type-specific epigenetic remodeling in prefrontal cortex begins before birth and continues into early adulthood. PLoS genetics 9, e1003433–e1003433, doi:10.1371/journal.pgen.1003433 (2013).

21 Heintzman, N. D. et al. Distinct and predictive chromatin signatures of transcriptional promoters and enhancers in the human genome. Nature genetics 39, 311–318, doi:10.1038/ng1966 (2007).

22 Marzi, S. J. et al. A histone acetylome-wide association study of Alzheimer’s disease identifies disease-associated H3K27ac differences in the entorhinal cortex. Nat Neurosci 21, 1618–1627, doi:10.1038/s41593-018-0253-7 (2018).

23 Kumar, A. et al. Age-associated changes in gene expression in human brain and isolated neurons. Neurobiology of aging 34, 1199–1209, doi:10.1016/j.neurobiolaging.2012.10.021 (2013).

24 Lu, T. et al. Gene regulation and DNA damage in the ageing human brain. Nature 429, 883–891, doi:10.1038/nature02661 (2004).

25 Khan, A. & Zhang, X. dbSUPER: a database of super-enhancers in mouse and human genome. Nucleic acids research 44, D164–171, doi:10.1093/nar/gkv1002 (2016).

26 Fu, Y., Rusznak, Z., Herculano-Houzel, S., Watson, C. & Paxinos, G. Cellular composition characterizing postnatal development and maturation of the mouse brain and spinal cord. Brain structure & function 218, 1337–1354, doi:10.1007/s00429-012-0462-x (2013).

27 Semple, B. D., Blomgren, K., Gimlin, K., Ferriero, D. M. & Noble-Haeusslein, L. J. Brain development in rodents and humans: Identifying benchmarks of maturation and vulnerability to injury across species. Progress in neurobiology 106-107, 1–16, doi:10.1016/j.pneurobio.2013.04.001 (2013).

28 Santos-Rosa, H. et al. Active genes are tri-methylated at K4 of histone H3. Nature 419, 407–411, doi:10.1038/nature01080 (2002).

29 Creyghton, M. P. et al. Histone H3K27ac separates active from poised enhancers and predicts developmental state. Proceedings of the National Academy of Sciences 107, 21931–21936, doi:10.1073/pnas.1016071107 (2010).

30 Handley, E. E. et al. Synapse Dysfunction of Layer V Pyramidal Neurons Precedes Neurodegeneration in a Mouse Model of TDP-43 Proteinopathies. Cerebral cortex (New York, N.Y. :1991) 27, 3630–3647, doi:10.1093/cercor/bhw185 (2017).

31 Ximerakis, M. et al. Single-cell transcriptomic profiling of the aging mouse brain. Nat Neurosci 22, 1696–1708, doi:10.1038/s41593-019-0491-3 (2019).

32 Swain, U. & Subba Rao, K. Study of DNA damage via the comet assay and base excision repair activities in rat brain neurons and astrocytes during aging. Mech Ageing Dev 132, 374–381, doi:10.1016/j.mad.2011.04.012 (2011).

33 Lodato, M. A. et al. Aging and neurodegeneration are associated with increased mutations in single human neurons. Science (New York, N.Y.) 359, 555–559, doi:10.1126/science.aao4426 (2018).

34 Suberbielle, E. et al. Physiologic brain activity causes DNA double-strand breaks in neurons, with exacerbation by amyloid-beta. Nat Neurosci 16, 613–621, doi:10.1038/nn.3356 (2013).

35 Madabhushi, R. et al. Activity-Induced DNA Breaks Govern the Expression of Neuronal Early-Response Genes. Cell 161, 1592–1605, doi:10.1016/j.cell.2015.05.032 (2015).

36 Yu, H. et al. Tet3 regulates synaptic transmission and homeostatic plasticity via DNA oxidation and repair. Nat Neurosci 18, 836–843, doi:10.1038/nn.4008 (2015).

37 Wu, W. et al. Neuronal enhancers are hotspots for DNA single-strand break repair. Nature 593, 440–444, doi:10.1038/s41586-021-03468-5 (2021).

38 Burke, S. N. & Barnes, C. A. Neural plasticity in the ageing brain. Nat Rev Neurosci 7, 30–40, doi:10.1038/nrn1809 (2006).

39 Toth, M. L. et al. Neurite Sprouting and Synapse Deterioration in the Aging *Caenorhabditis elegans* Nervous System. The Journal of Neuroscience 32, 8778, doi:10.1523/JNEUROSCI.1494-11.2012 (2012).

40 Mostany, R. et al. Altered synaptic dynamics during normal brain aging. J Neurosci 33, 4094–4104, doi:10.1523/JNEUROSCI.4825-12.2013 (2013).

41 Oh, G. et al. Epigenetic assimilation in the aging human brain. Genome Biology 17, 76, doi:10.1186/s13059-016-0946-8 (2016).

42 Douaud, G. et al. A common brain network links development, aging, and vulnerability to disease. Proceedings of the National Academy of Sciences 111, 17648, doi:10.1073/pnas.1410378111 (2014).

43 Blankvoort, S., Witter, M. P., Noonan, J., Cotney, J. & Kentros, C. Marked Diversity of Unique Cortical Enhancers Enables Neuron-Specific Tools by Enhancer-Driven Gene Expression. Curr Biol 28, 2103–2114 e2105, doi:10.1016/j.cub.2018.05.015 (2018).

44 de Magalhaes, J. P. Programmatic features of aging originating in development: aging mechanisms beyond molecular damage? FASEB J 26, 4821–4826, doi:10.1096/fj.12-210872 (2012).

45 Somel, M. et al. MicroRNA, mRNA, and protein expression link development and aging in human and macaque brain. Genome Res 20, 1207–1218, doi:10.1101/gr.106849.110 (2010).

46 Colantuoni, C. et al. Temporal dynamics and genetic control of transcription in the human prefrontal cortex. Nature 478, 519–523, doi:10.1038/nature10524 (2011).

47 Huang, B. et al. Inhibition of histone acetyltransferase GCN5 extends lifespan in both yeast and human cell lines. Aging Cell 19, e13129, doi:10.1111/acel.13129 (2020).

48 Kuo, Y. M. & Andrews, A. J. Quantitating the specificity and selectivity of Gcn5-mediated acetylation of histone H3. PLoS One 8, e54896, doi:10.1371/journal.pone.0054896 (2013).

49 Stilling, R. M. et al. K-Lysine acetyltransferase 2a regulates a hippocampal gene expression network linked to memory formation. EMBO J 33, 1912–1927, doi:10.15252/embj.201487870 (2014).

50 Bardai, F. H. & D’Mello, S. R. Selective toxicity by HDAC3 in neurons: regulation by Akt and GSK3beta. J Neurosci 31, 1746–1751, doi:10.1523/JNEUROSCI.5704-10.2011 (2011).

51 de Ruijter, A. J., van Gennip, A. H., Caron, H. N., Kemp, S. & van Kuilenburg, A. B. Histone deacetylases (HDACs): characterization of the classical HDAC family. Biochem J 370, 737–749, doi:10.1042/BJ20021321 (2003).

52 Tong, J. J., Liu, J., Bertos, N. R. & Yang, X. J. Identification of HDAC10, a novel class II human histone deacetylase containing a leucine-rich domain. Nucleic acids research 30, 1114–1123, doi:10.1093/nar/30.5.1114 (2002).

53 Nitarska, J. et al. A Functional Switch of NuRD Chromatin Remodeling Complex Subunits Regulates Mouse Cortical Development. Cell Rep 17, 1683–1698, doi:10.1016/j.celrep.2016.10.022 (2016).

54 Sokpor, G., Xie, Y., Rosenbusch, J. & Tuoc, T. Chromatin Remodeling BAF (SWI/SNF) Complexes in Neural Development and Disorders. Front Mol Neurosci 10, 243, doi:10.3389/fnmol.2017.00243 (2017).

55 Pisansky, M. T. et al. Mice lacking the chromodomain helicase DNA-binding 5 chromatin remodeler display autism-like characteristics. Transl Psychiatry 7, e1152, doi:10.1038/tp.2017.111 (2017).

56 Potts, R. C. et al. CHD5, a brain-specific paralog of Mi2 chromatin remodeling enzymes, regulates expression of neuronal genes. PLoS One 6, e24515, doi:10.1371/journal.pone.0024515 (2011).

57 Paredes, S. & Maggert, K. A. Ribosomal DNA contributes to global chromatin regulation. Proc Natl Acad Sci U S A 106, 17829–17834, doi:10.1073/pnas.0906811106 (2009).

58 Tiku, V. & Antebi, A. Nucleolar Function in Lifespan Regulation. Trends Cell Biol 28, 662–672, doi:10.1016/j.tcb.2018.03.007 (2018).

59 Srivastava, R., Srivastava, R. & Ahn, S. H. The Epigenetic Pathways to Ribosomal DNA Silencing. Microbiol Mol Biol Rev 80, 545–563, doi:10.1128/MMBR.00005-16 (2016).

60 Nelson, J. O., Watase, G. J., Warsinger-Pepe, N. & Yamashita, Y. M. Mechanisms of rDNA Copy Number Maintenance. Trends Genet 35, 734–742, doi:10.1016/j.tig.2019.07.006 (2019).

61 Hallgren, J., Pietrzak, M., Rempala, G., Nelson, P. T. & Hetman, M. Neurodegeneration-associated instability of ribosomal DNA. Biochim Biophys Acta 1842, 860–868, doi:10.1016/j.bbadis.2013.12.012 (2014).

62 Gaubatz, J., Prashad, N. & Cutler, R. G. Ribosomal RNA gene dosage as a function of tissue and age for mouse and human. Biochim Biophys Acta 418, 358–375, doi:10.1016/0005-2787(76)90297-5 (1976).

63 Oakford, P. C. et al. Transcriptional and epigenetic regulation of the GM-CSF promoter by RUNX1. Leuk Res 34, 1203–1213, doi:10.1016/j.leukres.2010.03.029 (2010).

64 Taberlay, P. C. et al. Polycomb-repressed genes have permissive enhancers that initiate reprogramming. Cell 147, 1283–1294, doi:10.1016/j.cell.2011.10.040 (2011).

65 Taberlay, P. C., Statham, A. L., Kelly, T. K., Clark, S. J. & Jones, P. A. Reconfiguration of nucleosome-depleted regions at distal regulatory elements accompanies DNA methylation of enhancers and insulators in cancer. Genome Res 24, 1421–1432, doi:10.1101/gr.163485.113 (2014).

66 Andrews, S. FastQC: A quality control tool for high throughput sequence data. Reference Source (2010).

67 Ewels, P., Magnusson, M., Lundin, S. & Kaller, M. MultiQC: summarize analysis results for multiple tools and samples in a single report. Bioinformatics (Oxford, England) 32, 3047–3048, doi:10.1093/bioinformatics/btw354 (2016).

68 Martin, M. Cutadapt removes adapter sequences from high-throughput sequencing reads. 2011 17, doi:10.14806/ej.17.1.200 pp. 10–12 (2011).

69 Langmead, B. & Salzberg, S. L. Fast gapped-read alignment with Bowtie 2. Nat Meth 9, 357–359, doi:10.1038/nmeth.1923 (2012).

70 Waterston, R. H. et al. Initial sequencing and comparative analysis of the mouse genome. Nature 420, 520–562, doi:10.1038/nature01262 (2002).

71 Diaz, A., Nellore, A. & Song, J. S. CHANCE: comprehensive software for quality control and validation of ChIP-seq data. Genome Biol 13, R98, doi:10.1186/gb-2012-13-10-r98 (2012).

72 Ramírez, F. et al. deepTools2: a next generation web server for deep-sequencing data analysis. Nucleic acids research 44, W160–W165, doi:10.1093/nar/gkw257 (2016).

73 Gjoneska, E. et al. Conserved epigenomic signals in mice and humans reveal immune basis of Alzheimer/’s disease. Nature 518, 365–369, doi:10.1038/nature14252 (2015).

74 Landt, S. G. et al. ChIP-seq guidelines and practices of the ENCODE and modENCODE consortia. Genome Res 22, 1813–1831, doi:10.1101/gr.136184.111 (2012).

75 Zhang, Y. et al. Model-based analysis of ChIP-Seq (MACS). Genome Biol 9, R137, doi:10.1186/gb-2008-9-9-r137 (2008).

76 Consortium, E. P. An integrated encyclopedia of DNA elements in the human genome. Nature 489, 57–74, doi:10.1038/nature11247 (2012).

77 Lun, A. T. L. & Smyth, G. K. csaw: a Bioconductor package for differential binding analysis of ChIP-seq data using sliding windows. Nucleic acids research 44, e45, doi:10.1093/nar/gkv1191 (2016).

78 Robinson, M. D., McCarthy, D. J. & Smyth, G. K. edgeR: a Bioconductor package for differential expression analysis of digital gene expression data. Bioinformatics (Oxford, England) 26, 139–140, doi:10.1093/bioinformatics/btp616 (2010).

79 Robinson, M. D. & Oshlack, A. A scaling normalization method for differential expression analysis of RNA-seq data. Genome Biology 11, R25, doi:10.1186/gb-2010-11-3-r25 (2010).

80 Ramirez, F. et al. deepTools2: a next generation web server for deep-sequencing data analysis. Nucleic acids research 44, W160–165, doi:10.1093/nar/gkw257 (2016).

81 McLean, C. Y. et al. GREAT improves functional interpretation of cis-regulatory regions. Nat Biotechnol 28, 495–501, doi:10.1038/nbt.1630 (2010).

